# Volatile molecules secreted by the wheat pathogen *Parastagonospora nodorum* are involved in development and phytotoxicity

**DOI:** 10.1101/729509

**Authors:** M. Jordi Muria-Gonzalez, Hui Yeng Yeannie Yap, Susan Breen, Oliver Mead, Chen Wang, Yit-Heng Chooi, Russell A. Barrow, Peter S. Solomon

## Abstract

Septoria nodorum blotch is a major disease of wheat caused by the fungus *Parastagonospora nodorum*. Recent studies have demonstrated that secondary metabolites, including polyketides and non-ribosomal peptides, produced by the pathogen play important roles in disease and development. However, there is currently no knowledge on the composition or biological activity of the volatile organic compounds (VOCs) secreted by *P. nodorum*. To address this, we undertook a series of growth and phytotoxicity assays and demonstrated that *P. nodorum* VOCs inhibited bacterial growth, were phytotoxic and suppressed self-growth. Mass spectrometry analysis revealed that 3-methyl-1-butanol, 2-methyl-1-butanol, 2-methyl-1-propanol and 2-phenylethanol were dominant in the VOC mixture and phenotypic assays using these short chain alcohols confirmed that they were phytotoxic. Further analysis of the VOCs also identified the presence of multiple sesquiterpenes of which four were identified via mass spectrometry and nuclear magnetic resonance as β-elemene, α-cyperone, eudesma-4,11-diene and acora-4,9-diene. Subsequent reverse genetics studies were able to link these molecules to corresponding sesquiterpene synthases in the *P. nodorum* genome. However, despite extensive testing, these molecules were not involved in either of the growth inhibition or phytotoxicity phenotypes previously observed. Plant assays using mutants of the pathogen lacking the synthetic genes revealed that the identified sesquiterpenes were not required for disease formation on wheat leaves. Collectively, these data have significantly extended our knowledge of the VOCs in fungi and provided the basis for further dissecting the roles of sesquiterpenes in plant disease.

## Introduction

The Dothideomycete fungus *Parastagonospora nodorum* is the causal agent of Septoria nodorum blotch, a significant foliar global disease of wheat. Once considered a simplistic pathogen that caused disease through the secretion of lytic enzymes, seminal studies over the last decade have demonstrated that *P. nodorum* facilitates disease through the use of small proteins called effectors ^1^. To date, three effectors from *P. nodorum* have been described, ToxA, Tox1 and Tox3 ^2–4^. Each of these proteins interacts in a gene-for-gene for manner with specific cognate susceptibility genes in the host leading to host cell death and disease. More recent studies have demonstrated that as well as inducing necrosis, each of these effectors appears to also function in repressing host defence responses highlighting the complex nature of this interaction ^5–7^.

However, it has been recently shown that ToxA, Tox1 and Tox3 are not the only molecules responsible for *P. nodorum* to successfully infect wheat ^8,9^. Tan *et al.* (2015) used a reverse genetic approach to generate a strain of *P. nodorum* lacking each of the effector genes and showed that the resulting mutant, albeit being less pathogenic, retained the ability to cause disease^8^. Indeed, recent studies have examined the role of several polyketide secondary metabolites synthesized by *P. nodorum* and shown that some have a role in facilitating disease on wheat ^10–13^. However, there are many more secondary metabolites encoded for within the *P. nodorum* genome that potentially play a role in the interaction of the pathogen with its host ^14,15^.

Another group of molecules that have yet to be characterised in terms of their role or requirement in septoria tritici blotch are the volatile organic compounds (VOCs). VOCs are small carbon-based molecules that readily evaporate and are ubiquitously produced by most forms of life ^16^. It has been proposed that VOCs play important roles as signals in inter and intra-organismic interactions which surpasses the involvement of other diffusible molecules^17^. Microorganisms are known to be a rich source of VOCs displaying antibacterial, antifungal and phytotoxic properties, but also acting as chemical cues that help structuring microbial communities ^17^. VOCs typically produced by microorganisms are complex blends of chemical. The composition and role of VOCs though in fungi, particularly plant pathogens, are poorly understood.

To address this knowledge gap, we firstly explored the biological activity of the VOCs emitted by *P. nodorum* and undertook an initial identification and characterisation of the major components. As a result of this, several sesquiterpene molecules were identified and the genes required for their synthesis characterised. This study has shed further light on the chemical diversity synthesized by these fungi and has raised further questions as to the roles of small molecules generated by this devastating wheat pathogen.

## Results

### *Parastagonospora nodorum* VOCs have phytotoxic, antibiotic and self-inhibitory properties

To assess if volatile emissions of *P. nodorum* harbour bioactive VOCs, split plate assays were performed to evaluate a series of biological activities including phytotoxicity, fungitoxicity and bactericidal (Figure 1). Segmented Petri dishes were used to prevent molecules diffusing through the media and ensure that any observable activities could be solely attributed to the volatile compounds. To assay for phytotoxicity, seeds from the host of *P. nodorum*, wheat *cv*. Grandin, and a dicotyledonous plant, *Medicago truncatula*, were used. *P. nodorum* VOCs had a strong effect on the radicle elongation and hypocotyl or coleoptile growth which were significantly reduced when the seeds were germinated in the presence of *P. nodorum* but appeared unaffected in the control plates (no fungal inoculation). The effect on bacteria was mixed as there was no observable impact on the growth of a variety of different strains including *Escherichia coli, Pseudomonas syringae, Bacillus cereus*, or *Flavobacterium* sp. in the presence of the fungal VOCs (Figure S1). In contrast, there was a strong reduction in the growth of the nitrogen-fixing bacterium *Sinorhizobium meliloti* and also *Sphingobacterium multivorum* when *P. nodorum* was cultured in the same Petri dish (Figure 1). There was no apparent impact on the growth of any of the fungi tested when grown with *P. nodorum* with the exception of the apparent self-inhibition of *P. nodorum* growth (by its own VOCs) (Figure 1, Figure S1).

**Figure 1.**
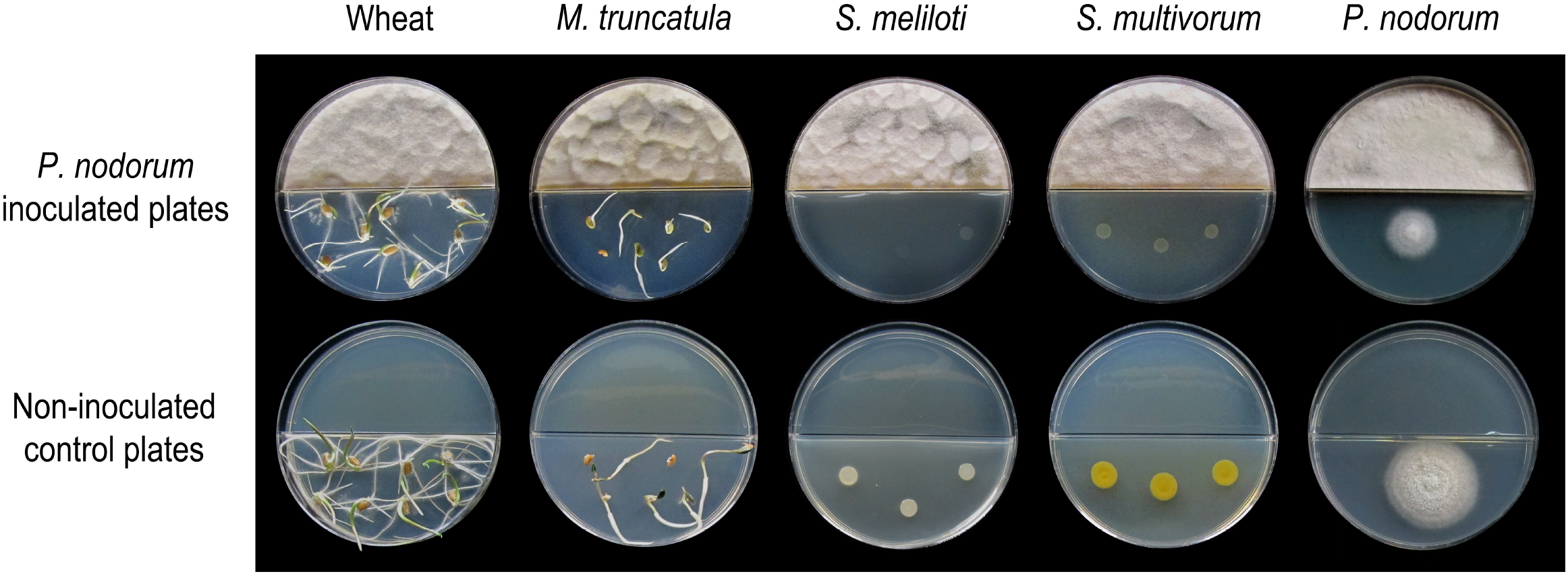
Split plate assays of the effect of VOCs produced by wild type *P. nodorum* (SN15). The top row of plates show the inhibition caused by *P. nodorum* VOCs on wheat and *Medicago* seeds, *S. meliloti, S. multivorum* and *P. nodorum*. The bottom row of plates shows the growth of each of the test organisms in the absence of *P. nodorum*.

### The major *Parastagonospora nodorum* VOCs are short chain alcohols

To dissect the chemical basis of the bioactivities described above, the identities of the volatile molecules were determined using a combination of solid phase micro-extractions (SPME) from the headspace (HS) of ten days old fungal cultures in slanted Fries agar vials and subsequent analysis by gas chromatography-mass spectrometry (GC-MS) and spectral comparison against pure standard and the NIST library. Within the *P. nodorum* VOCs mixture, several alcohols and esters were identified as being the most prominent signals (percentage of area of the whole chromatogram) (Table 1): 3-methyl-1-butanol (representing 5.36% of the VOCs mixture), 2-methyl-1-butanol (2.6%), 2-methyl-1-propanol (1.43%) and 2-phenylethanol (1.13%). Many other volatile molecules were also identified in peaks with smaller areas. The polyketide mellein (0.9%) was also detected along with some sesquiterpenes of which two were putatively identified as β-elemene and eudesma-4,11-diene.

**Table 1.**
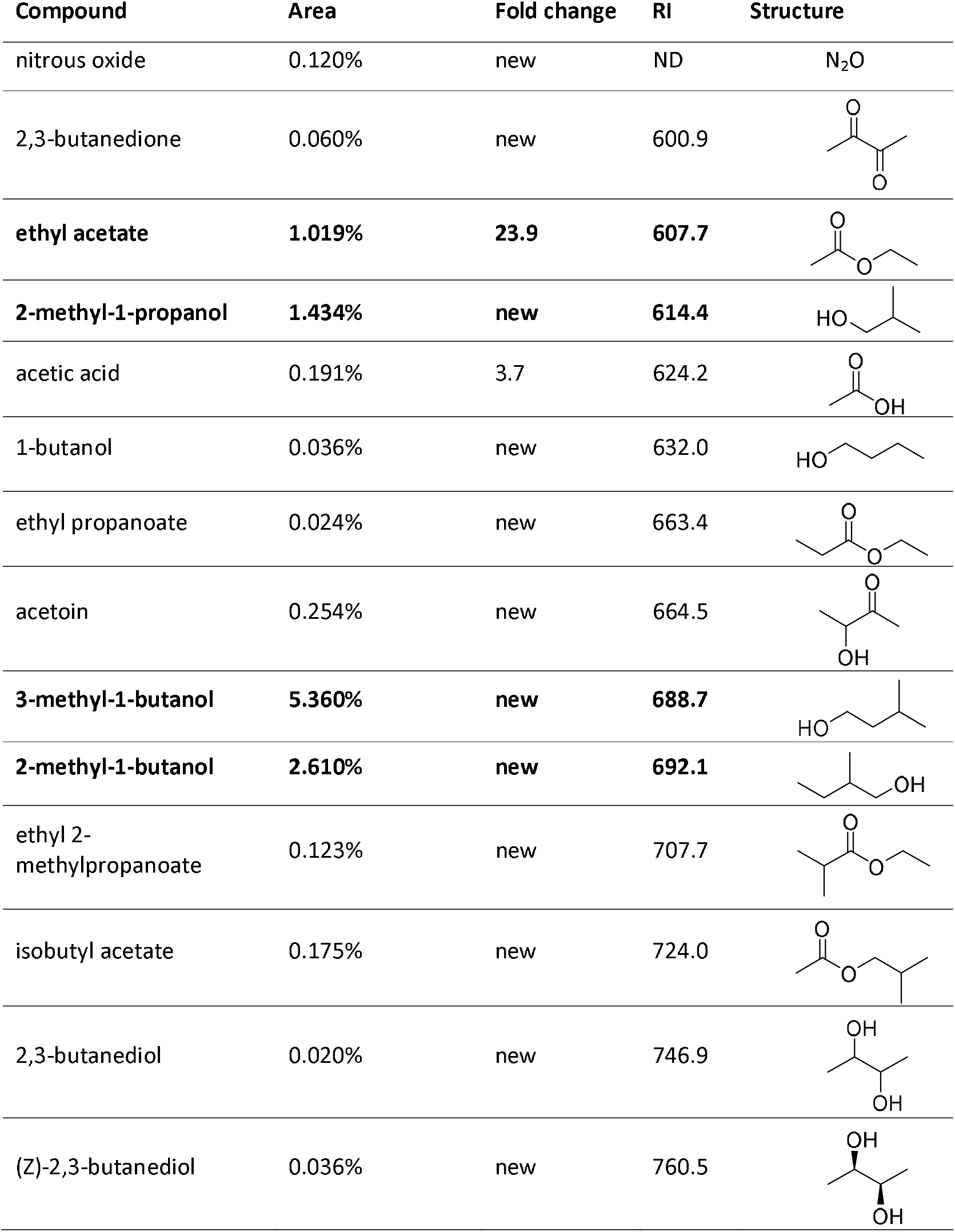
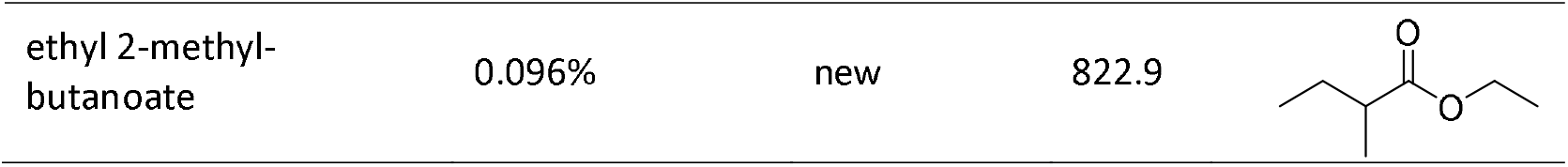
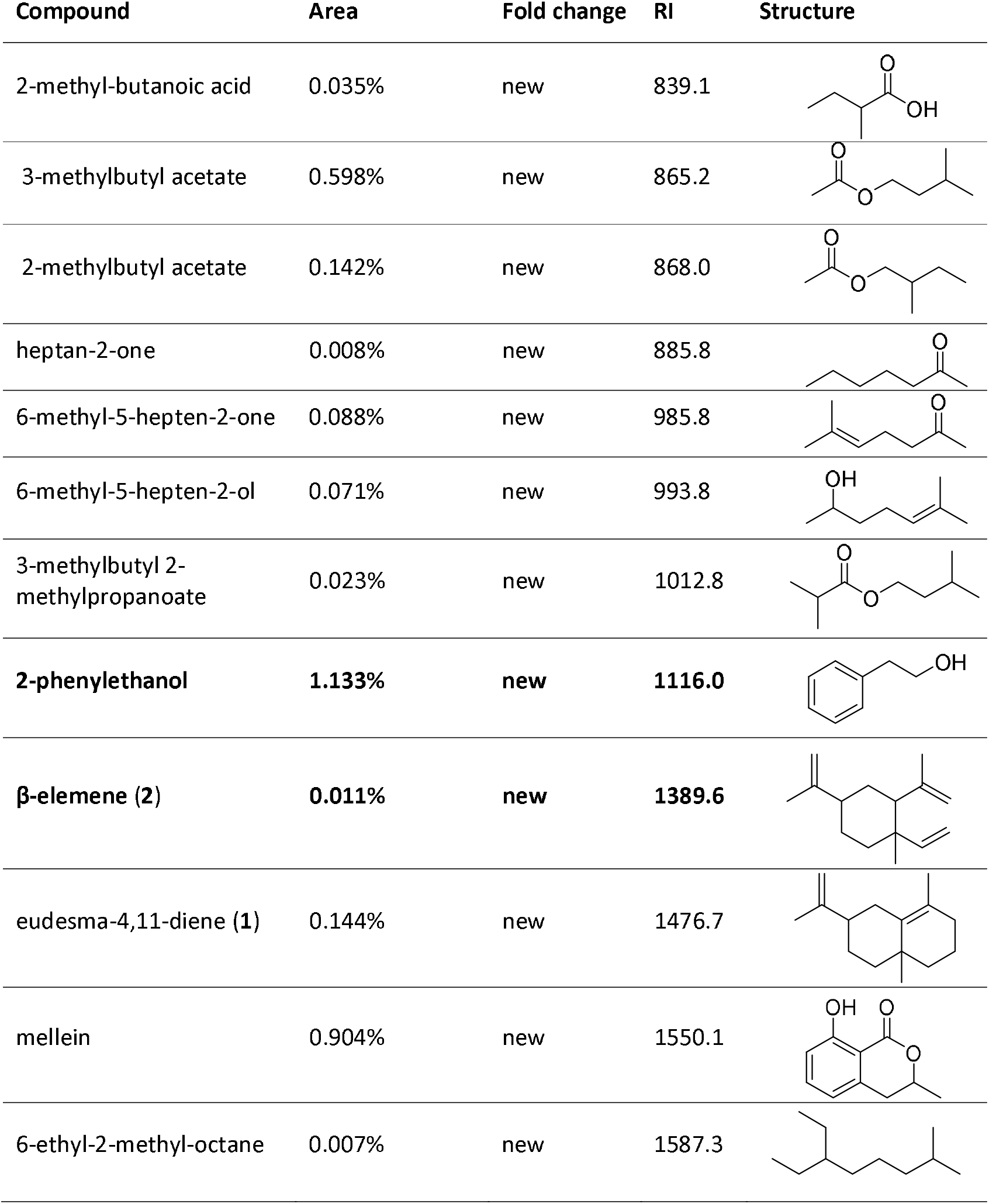
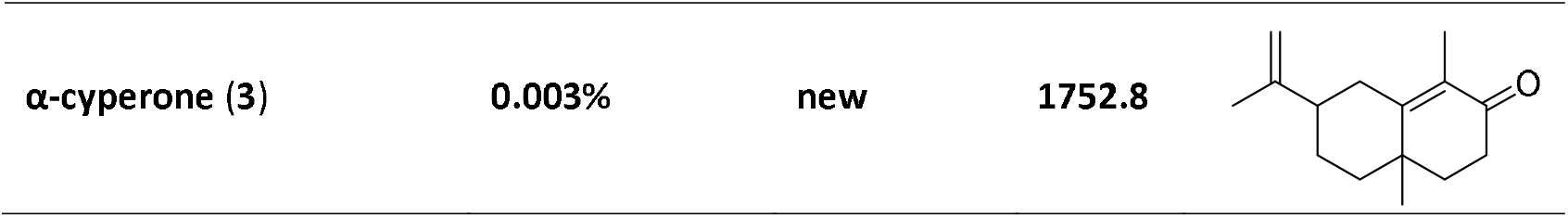
VOCs identified from *P. nodorum* grown on Fries media using HS-SPME-GC-MS. Compound identification was done by comparison of the mass spectra against the NIST library and, for those in bold, against standards. In all cases stereochemistry was not determined. Area indicates the percentage of a particular peak’s area compared to the total chromatogram’s area under the curve.

| Compound | Area | Fold change | RI | Structure |
| --- | --- | --- | --- | --- |
| nitrous oxide | 0.120% | new | ND | N<sub>2</sub>O |
| 2,3-butanedione | 0.060% | new | 600.9 |  |
| **ethyl acetate** | **1.019%** | **23.9** | **607.7** |  |
| **2-methyl-1-propanol** | **1.434%** | **new** | **614.4** |  |
| acetic acid | 0.191% | 3.7 | 624.2 |  |
| 1-butanol | 0.036% | new | 632.0 |  |
| ethyl propanoate | 0.024% | new | 663.4 |  |
| acetoin | 0.254% | new | 664.5 |  |
| **3-methyl-1-butanol** | **5.360%** | **new** | **688.7** |  |
| **2-methyl-1-butanol** | **2.610%** | **new** | **692.1** |  |
| ethyl 2-methylpropanoate | 0.123% | new | 707.7 |  |
| isobutyl acetate | 0.175% | new | 724.0 |  |
| 2,3-butanediol | 0.020% | new | 746.9 |  |
| (Z)-2,3-butanediol | 0.036% | new | 760.5 |  |
| ethyl 2-methyl-butanoate | 0.096% | new | 822.9 |  |

**Table 1** (continued). VOCs identified from *P.
| Compound | Area | Fold change | RI | Structure |
| --- | --- | --- | --- | --- |
| 2-methyl-butanoic acid | 0.035% | new | 839.1 |  |
| 3-methylbutyl acetate | 0.598% | new | 865.2 |  |
| 2-methylbutyl acetate | 0.142% | new | 868.0 |  |
| heptan-2-one | 0.008% | new | 885.8 |  |
| 6-methyl-5-hepten-2-one | 0.088% | new | 985.8 |  |
| 6-methyl-5-hepten-2-ol | 0.071% | new | 993.8 |  |
| 3-methylbutyl 2-methylpropanoate | 0.023% | new | 1012.8 |  |
| **2-phenylethanol** | **1.133%** | **new** | **1116.0** |  |
| **β-elemene (2)** | **0.011%** | **new** | **1389.6** |  |
| eudesma-4,11-diene (1) | 0.144% | new | 1476.7 |  |
| mellein | 0.904% | new | 1550.1 |  |
| 6-ethyl-2-methyl-octane | 0.007% | new | 1587.3 |  |
| **α-cyperone (3)** | **0.003%** | **new** | **1752.8** |  |

### The four most abundant *Parastagonospora nodorum* VOCs are phytotoxic

The activity of the four most prominent volatile molecules identified in the chromatograms from the head space of *P. nodorum* (3-methyl-1-butanol, 2-methyl-1-butanol, 2-methyl-1-propanol, 2-phenylethanol) were assayed to assess their impact on the growth of *P. nodorum* and wheat seedling development. These compounds were tested independently at an atmospheric concentration of 1 mM. Additionally, a mixture of these compounds following the *in vitro* proportions, was prepared and tested at 100 ppm. Neither the independent pure VOCs nor the mixture had any effect on *P. nodorum* growth suggesting that these molecules are not responsible for the inhibitory effect described above (data not shown). In contrast, a 27% decrease in germination was observed when wheat seeds were exposed to 3-methyl-1-butanol, although no other treatment had a significant effect on germination (Figure 2). However, both radicle and coleoptile elongation were repressed in all treatments; 3-methyl-1-butanol showed the greatest inhibition (100% and 83% respectively) while 2-methyl-1-propanol showed the least inhibition (36% and 31% respectively).

**Figure 2.**
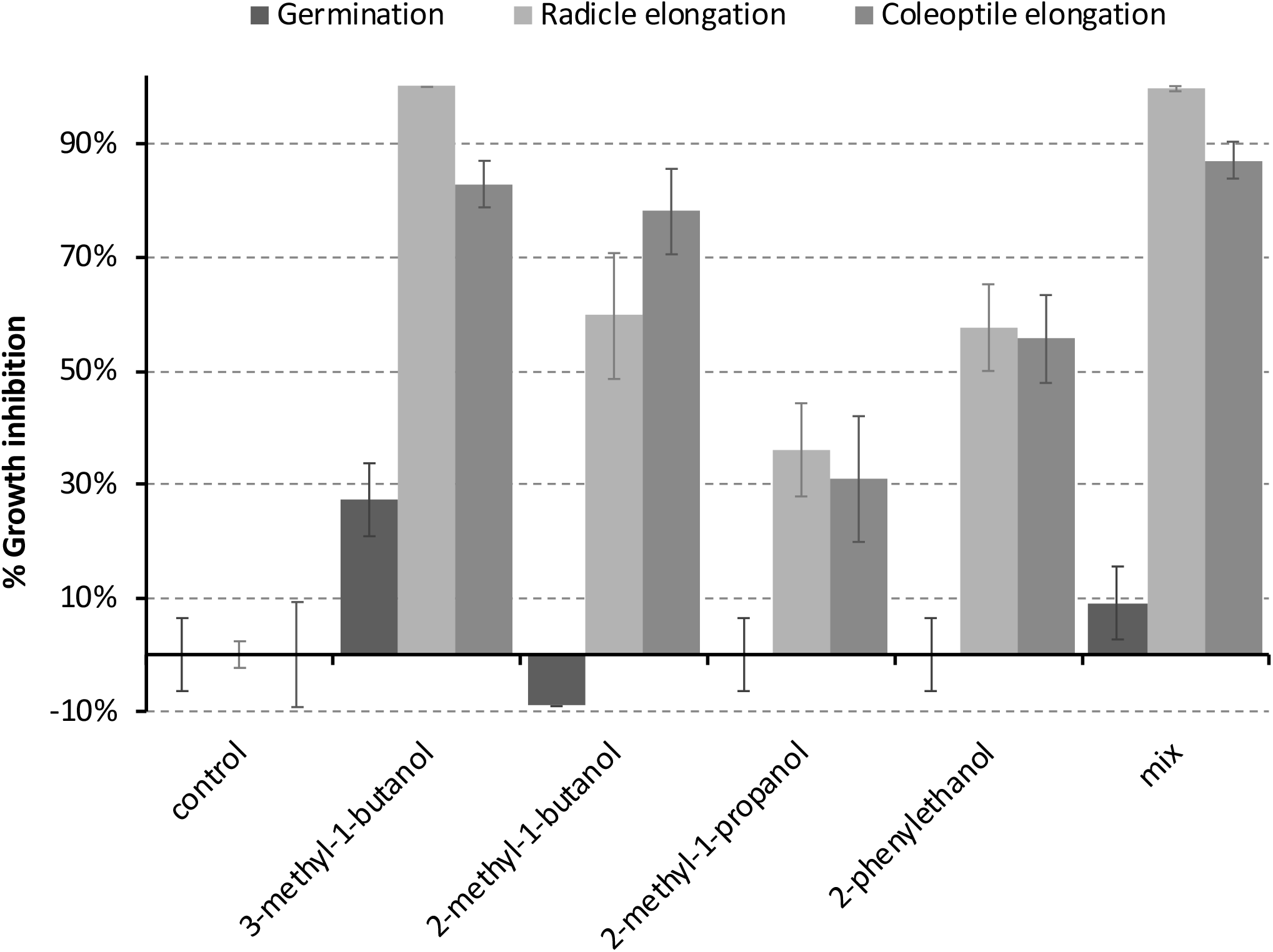
Inhibition of wheat seedling development by the predominant VOCs produced by *P. nodorum*. The bars represents the inhibitory activity on the developmental stage. 3-methyl-1-butanol, 2-methyl-1-butanol, 2-methyl-1-propanol and 2-phenylethanol were tested at 1 mM. Error bars show the standard deviation.

### *In planta* production of sesquiterpenes

In addition to the bioactive short chain alcohols, we were also interested in the presence of the sesquiterpenes found in the axenic culture VOCs To determine if these molecules played a potential role in disease development, the production of sesquiterpenes was assayed for during infection and compared to those produced in axenic culture. VOCs were extracted from vials containing either infected leaves or the fungus grown axenically and analysed by HS-SPME-GC-MS. Interestingly the same sesquiterpenes produced *in vitro* by *P. nodorum* were also found *in planta* as well as others not previously observed (Figure 3). Eudesma-4,11-diene (sesquiterpene **1**), β-elemene (sesquiterpene **2**) and α-cyperone (sesquiterpene **3**) were putatively identified by comparing the acquired data against the NIST database. Another interesting compound was sesquiterpene **4**, the most abundant sesquiterpene detected from *P. nodorum*, in Fries cultures and in wheat leaves. However, despite a fragmentation pattern and molecular weight typical of a sesquiterpene, its match against entries present in the NIST library was not high enough to confidently assign a possible identity.

**Figure 3.**
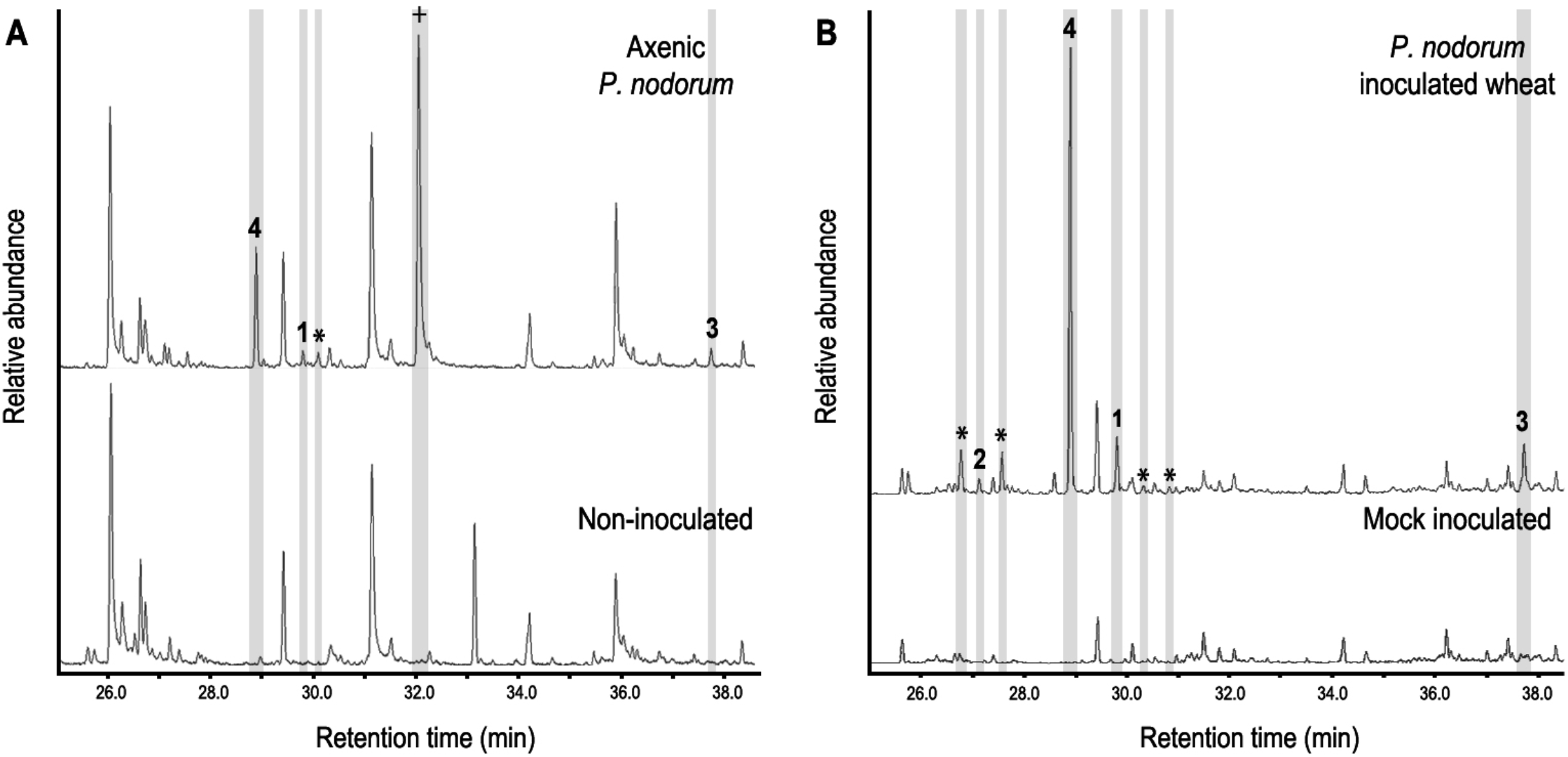
(A) Sesquiterpene profile of *P. nodorum* growing on Fries media (upper panel) and uninoculated Fries media (lower panel). (**B**) The sesquiterpene profile of wheat leaves infected with *P. nodorum* (upper panel) and mock inoculated wheat leaves (low panel). Compounds **1-4** are shown and other sesquiterpenes peaks are stared (*). Mellein is indicated as a cross (+).

### The biosynthetic genes of *P. nodorum* sesquiterpenes

The *P. nodorum* genome encodes three sesquiterpene synthases ^14^, *Sts1*, *Sts2* and *Sts3*. Previous studies have demonstrated that *Sts1* and *2* are expressed during infection, but not *Sts3*. Furthermore, analysis of the *Sts3* gene sequence revealed that it appears truncated implying that it isn’t functional and thus it wasn’t considered for further study ^18^. To directly link the molecules identified above to the genes, *Sts1* and *Sts2* were disrupted individually in the *P. nodorum* genome through homologous recombination. Disruption cassettes were constructed to independently replace *Sts1* and *Sts2* with a phleomycin resistance marker. *P. nodorum* was transformed and positive colonies were selected from phleomycin-containing plates. Correct disruption of the genes was verified by PCR and strains containing a single copy of the disruption cassette were selected by qPCR.

Single copy transformants of *P. nodorum* strains lacking *Sts1* (sts1) and *Sts2* (sts2) were selected for further analysis by HS-SPME-GC-MS (Figure 4). The signals of sesquiterpenes **1**, **2** and **3** along with three other unidentified sesquiterpenes present in wild-type *P. nodorum* were absent in the sts2 mutants indicating this gene codes for the core biosynthetic enzyme of the three putative sesquiterpenes plus some other sesquiterpene structures (same mass and similar fragmentation pattern). The mutants lacking *sts1* were missing three unidentified compounds putatively identified as sesquiterpenes, including sesquiterpene **4**, suggesting that *Sts1* is responsible for its synthesis.

**Figure 4.**
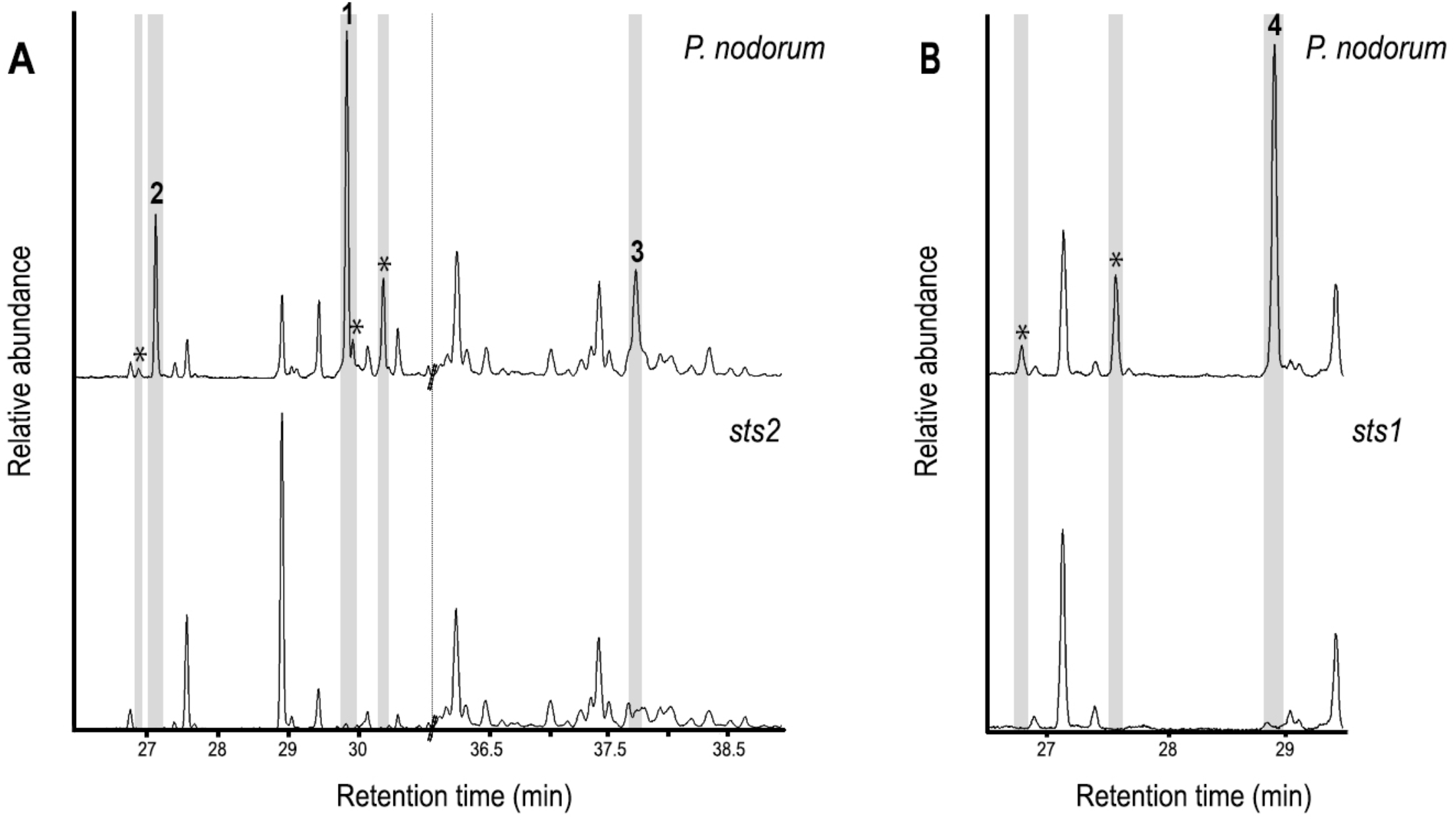
GC-MS chromatograms comparing sesquiterpenes in the extracts from wild-type *P. nodorum* to mutant strains lacking Sts2 (A) and Sts1 (B). The presence of **1 – 4** in the wild-type strain is highlighted.

### Heterologous expression of *Sts1* and *Sts2* reveals its prolificity and allows the sesquiterpenes isolation

To confirm the identity of these sequiterpenes, an isolation from medium scale fermentation of *P. nodorum* in Fries media was undertaken. Sesquiterpene **3** was isolated by silica flash chromatography followed by C18 flash chromatography. However, the isolation of sesquiterpenes **1**, **2** and **4** was not achieved. Consequently, the *Sts1* and *Sts2* genes were heterologously expressed in yeast to confirm **1** and **2** as eudesma-4,11-diene and β–elemene respectively. Acetone extracts from small scale cultures of the yeast strains harbouring the sesquiterpene synthases were analysed by GC-MS and the production of **1**, **2** and **4** by the heterologous expression of *Sts2* and *Sts1* was confirmed (Figure 5). Interestingly, sesquiterpene **3** (putatively α-cyperone) was not detected, suggesting it is modified post synthesis.

**Figure 5.**
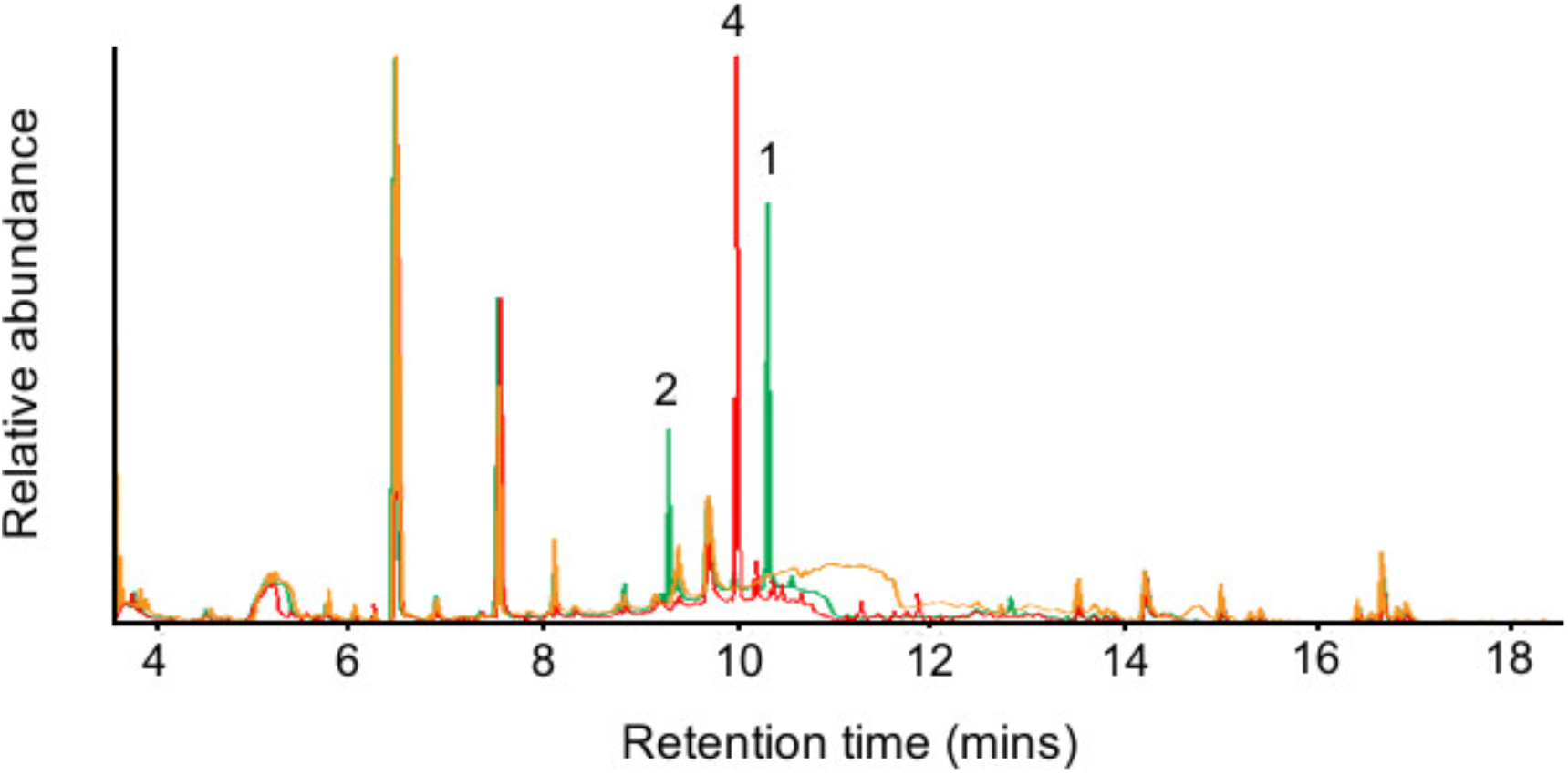
GC-MS chromatograms of the extracts from the *Sts1* and *Sts2* heterologous expression in yeast. Overlapped chromatograms of extracts from yeast harbouring *Sts1* (red), *Sts2* (green) and an empty vector (orange). The presence of **1, 2** and **4** is indicated.

Sesquiterpenes **1** and **2**, and **4** were isolated by silica followed by C18 flash chromatography of acetone extracts from medium scale YPDA fermentations of yeast carrying *Sts2* and *Sts1* respectively. Subsequent GC-MS analysis of the sesquiterpene fractions confirmed that **4** is the major product of Sts1 but also that 14 other unidentified terpenes are synthesised by the same enzyme (Figure S2). Similarly, the predominant sesquiterpene produced by Sts2 is **1** along with 10 other molecules including compound **2** (Figure S3).

### The identity of the *P. nodorum* sesquiterpenes are confirmed by MS^2^ and NMR

Commercial standards of β-elemene and α-cyperone were purchased and analysed by GC-MS^2^ to confirm the identities of sesquiterpenes **2** and **3**. Identical retention times in addition to a comparison of the MS^2^ fragmentation profiles that demonstrated a complete overlap of the major ions present in the standards compared to the extracted samples confirmed the identities of **2** and **3** (Figure 6, S4 and S5).

**Figure 6.**
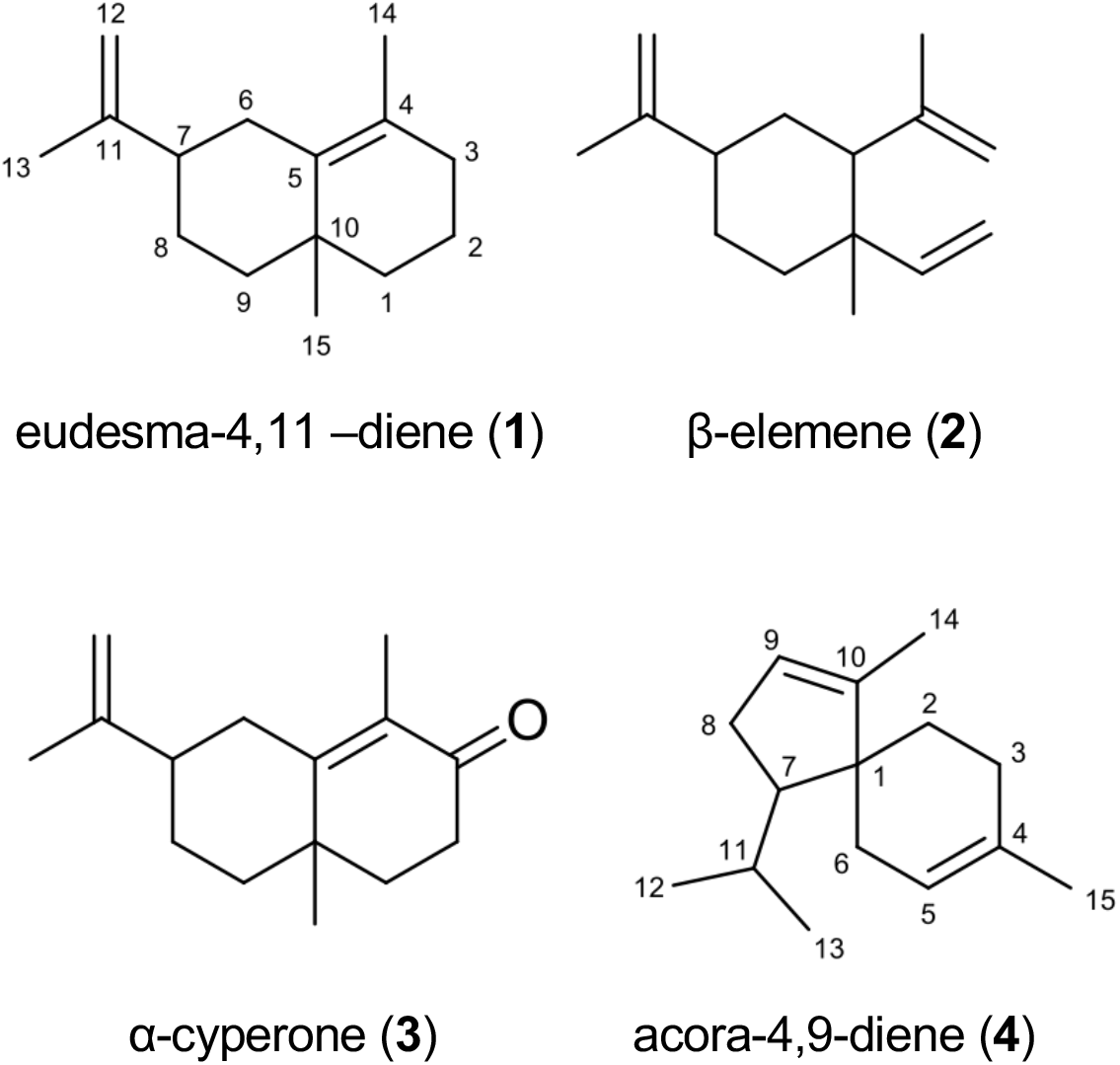
Identified sesquiterpenes present in *P. nodorum* VOCs

The isolated sesquiterpene **1**, the putative eudesma-4,1-diene, and unknown sesquiterpene **4** were subjected to ^1^H NMR, ^13^C NMR, HMBC and HSQC experiments (Figures S6 to S13). Tables S3 and S4 present the assignments and chemical shifts (δ) for all carbons and the corresponding hydrogens as well as the correlations obtained from the HMBC experiment for sesquiterpene **1** and sesquiterpene **4** respectively. The identity of sesquiterpene **1** was confirmed to be eudesma-4,11-diene, and sesquiterpene **4** is proposed to be acora-4,9-diene (Figure 6) based on comparison to previously reported NMR data ^19,20^.

### *Sts1* and *Sts2* are not required for bacterial growth inhibition, phytotoxicity or for the infection of wheat by *P. nodorum*

A segmented plate growth assay was used to determine if the molecules derived from either *Sts1* or *Sts2* are involved in the biological activities described above. As previously observed, volatiles from wild type *P. nodorum* reduced wheat seed germination and inhibited the growth of *S. multivorum* as well as self-growth. The growth of both the *sts1* or *sts2* mutants caused the same phenotypic effects as the wild type inferring that the products of either gene are not responsible for these inhibitory activities (data not shown).

The role of the Sts1 and Sts2 terpenes in pathogenicity was evaluated by inoculating the second leaf of two-week-old wheat seedlings with wild type *P. nodorum* and each of the mutants. Symptom development for the *sts1* and *sts2* mutants was unaffected compared to the wild type suggesting that the genes do not play a role in disease as assayed in this attached leaf system (Figure S14).

## Discussion

The roles and functions of volatile organic compounds produced by fungi are poorly understood. As such, we embarked upon a study to determine if the VOCs produced by the wheat pathogen *P. nodorum* were biologically active, and if so, resolve their identity. In this study, we have demonstrated that the wheat pathogen *P. nodorum* produces a range of VOCs that harbour intrinsic biological activities including inhibiting effects on plant seedlings, bacterial growth and also self-growth.

Initial assays clearly displayed that *P. nodorum* secretes bioactive VOCs as observed through the growth inhibition and phytotoxicity. It was also interesting to see evidence that the secreted VOCs may have a role in self growth regulation of the pathogen. Previous studies have demonstrated that volatiles can function in fungal self-inhibition of growth. For example, 1-octen-3-ol, a short chain alcohol produced by *Penicillium paneum*, and the sesquiterpene thujopsene from *Penicillium decumbes*, are known to inhibit the growth of the source fungi ^21,22^. Similarly, the selective inhibition of the *P. nodorum* VOC complement against bacteria observed is not without precedent. It has been previously demonstrated that compounds produced by sponge-associated Arctic microbial communities show a strong inhibitory activity against the opportunistic pathogenic bacterial *Burkholderia cepacia* complex but not to other pathogenic bacteria ^23,24^. Why VOCs from *P. nodorum* would harbour this specificity is unclear but it has been suggested that such selectivity may be a reflection of how different organisms respond differently to the same chemical cue or alternatively it may be a consequence on possible fitness differences among individuals. Such an effect of VOCs on shaping microbial communities has been previously proposed ^25^.

An analysis of the *P. nodorum* VOC chromatograms revealed that the four most prominent compounds are well-described short chain alcohols (Table 1). 3-Methyl-1-butanol and 2-methyl-1-butanol have been previously demonstrated to inhibit the growth of the fungal pathogen *Sclerotinia sclerotiorum* ^26^. Similarly, 2-phenylethanol affects gene expression and interferes in epigenetic regulation leading to the growth inhibition of multiple fungi including *Aspergillus flavus, Neurospora crassa* and *Penicillium spp*. ^27–29^. In contrast, low concentrations of 2-phenylethanol stimulates, rather than inhibits, the growth of *A. flavus*, revealing a hormetic behaviour of this compound ^30^. These data may suggest the involvement of these short chain alcohols in metabolic regulation and could be generalised communication signals in fungi. Furthermore, considering that these molecules are produced by a broad range of organisms, one could hypothesise that communication occurs at various levels ranging from interspecific to inter-kingdom crosstalk. Interestingly, when tested in pure form or artificially blended, these VOCs had no observable effect on the development of *P. nodorum*.

The effect of the four main *P. nodorum* volatile alcohols over plant and bacterial development has also been previously described. Akin to our observations, VOCs from truffles (*Tuber spp*.) inhibited the development of *Arabidopsis thaliana* ^31^. Within the volatile emissions of tuber fruiting bodies, 3-methyl-1-butanol inhibited *A. thaliana* germination at 130 ppm while 2-phenylethanol was inhibitory at 13 ppm and caused discoloration of the cotyledons of germinated seedlings at 130 ppm ^31^.

In contrast, VOCs mixtures produced by rhizobacteria containing 3-methyl-1-butanol, 2-methyl-1-butanol and 2-methyl-1-propanol promote growth of *A. thaliana* ^32,33^. It is possible that this differential effect is caused by variations in the proportion of the components of the volatilomes. It is known that differences in VOC levels in the soil correlate to changes in microbial soil populations ^34^. If we consider that interaction between organisms is a biological network that interweaves at different levels ^35^, it could be speculated that common volatile metabolites may help to coordinate the network by allowing organisms to eavesdrop on the communication signals from their neighbours ^36–38^.

Together with the short chain alcohols, a suite of sesquiterpenes were also identified in the VOC mixture. The presences of eudesma-4,11-diene (**1**), β-elemene (**2**), α-cyperone (**3**), and acora-4,9-diene (**4**) were all confirmed through a combination of mass spectrometry and NMR analysis. Subsequent reverse genetics and overexpression experiments then confirmed that the sesquiterpene synthase genes in *P. nodorum*, *Sts1* and *Sts2*, were responsible for their biosynthesis. Given the presence and abundance of these molecules during infection of wheat by the pathogen, it was surprising that mutant strains of the fungus lacking these molecules appeared unaffected in terms of development or pathogenicity. Indeed, information is scarce on what precisely the functions of these sesquiterpenes identified above are.

Many plants emit eudesma-4,11-diene (**1**) as a minor component of biologically active VOCs mixtures or essential oils ^39–43^. While **1** is produced by some basidiomycete and ascomycete fungi and by some actinomycetes, no biological activity has been described for either fungi or bacteria ^44–48^. Furthermore, **1** is produced (along with other sesquiterpenes) by soldier termites from different species and *Ceroplastes ceriferu*, a scale insect ^49–51^. Similarly, many plants and insects such as termites, aphids, butterflies and lady beetles, produce β-elemene (**2**) ^52–58^. The ascomycetes *Penicillium clavigerum, Penicillium roqueforti* and an endophytic *Nodulisporium* sp., and the basidiomycetes *Inonotus obliquus* and *Piptoporus betulinus*, are known to produce β-elemene ^44,47,59–61^. However, no role for this molecule in fungi has been identified.

The distribution of α-cyperone (**3**) seems to be more restricted. The molecule was first identified from the rhizomes of *Cyperus rotundus*, a medicinal plant which is also classified as an invasive grass ^62^. It has been postulated that **3** is causal to the described antimicrobial, phytotoxic, insecticidal, anti-inflammatory and antimalarial activities in essential oils from *C. rotundus*. ^63–66^. In fungi there is just one report corresponding to a stereoisomer of **3** isolated from an *endophytic Ascochyta sp*. from *Meliotus dentatus* but it has the opposite configuration to the plant isolated α-cyperone ^67^. Importantly, there are no reports on what this change in stereochemistry has on the function of the molecule. *Ascochyta* and *P. nodorum* are closely related fungi so it would not be unexpected if the α-cyperone identified in this study was also the opposite stereochemistry to the plant-derived molecule. As such, it is difficult to infer what the function of α-cyperone in *P. nodorum* may be.

In contrast to the widespread occurrence of molecules **1-3** discussed above, acora-4,9-diene (**4**), synthesised by *Sts1* in *P. nodorum* has only been found in the oils of vetiver (*Chrysopogon zizanioides*), in the seeds of carrot (*Daucus carota*), and in the glandular trichome exudates from leaves of Japanese rose (*Rosa rugosa*), with no biological activity described up to date ^68–70^.

Proposing an ecological role for Sts1 and Sts2 sesquiterpenes in the *P. nodorum*-wheat pathosystem is difficult due to the diversity of producers and reported activities of eudesma-4,11-diene (**1**) and β-elemene (**2**), the uncertainty of the *P. nodorum* α-cyperone (**3**) stereochemistry, the lack of information about acora-4,9-diene (**4**), and the absence of evident effects on pathogenicity, phytotoxicity, antimicrobial or self-regulating properties. Nonetheless, the *in planta* production of these molecules suggests its possible involvement in the fungal-plant interaction. Subtle changes in the interaction conferring some “competitive” advantage to the pathogen may not be easily detected in the laboratory pathogenicity tests which are not indicative of the full disease cycle of the pathogen.

Aside from the ecological role of the sesquiterpenes, the linking of genes to products in this study also provides an opportunity to better understand the biosynthesis of the identified sesquiterpenes. A non-redundant BlastP analysis of Sts1 and Sts2 suggested that the two proteins are related to trichodiene and aristolochene synthases respectively (Figure S15). These two enzymes have many similarities; in both cases the linear precursor of their products is farnesyl pyrophosphate (FPP) which loses its phosphate group and cyclises into a cationic cyclic intermediate. The difference is the type of carbocycle produced, which depends on the tertiary structure of the enzyme. While trichodiene synthase produces a bisabolyl cation, the aristolochene synthase produces a germacranyl cation ^71^. The sequence similarity of Sts1 and trichodiene synthases is congruent with the fact that the structure of acora-4,9-diene, and the other 14 putative sesquiterpenes produced by Sts1, seems to require a bisabolene intermediate (Figure 7). Conversely, a germacranyl intermediate is the likely intermediary of eudesma-4,11-diene, β-elemene, and the other 10 products of Sts2, which corresponds to the similarity between this enzyme and aristolochene synthases (Figure 7). The generation of the multiple products by a single sesquiterpene synthase is due to the intermediaries, bisabolyl and germacranyl cations in this case, suffering spontaneous rearrangements with minimal involvement of the biosynthetic enzyme. Terpene cyclases displaying a higher control over these subsequent reactions may produce fewer structures or even a single product ^72^. The generation of a wide spectrum of sesquiterpenes or other chemical structures increase the chances of some of these molecules having the right conformation to interact with diverse biological targets and affecting other organisms

**Figure 7.**
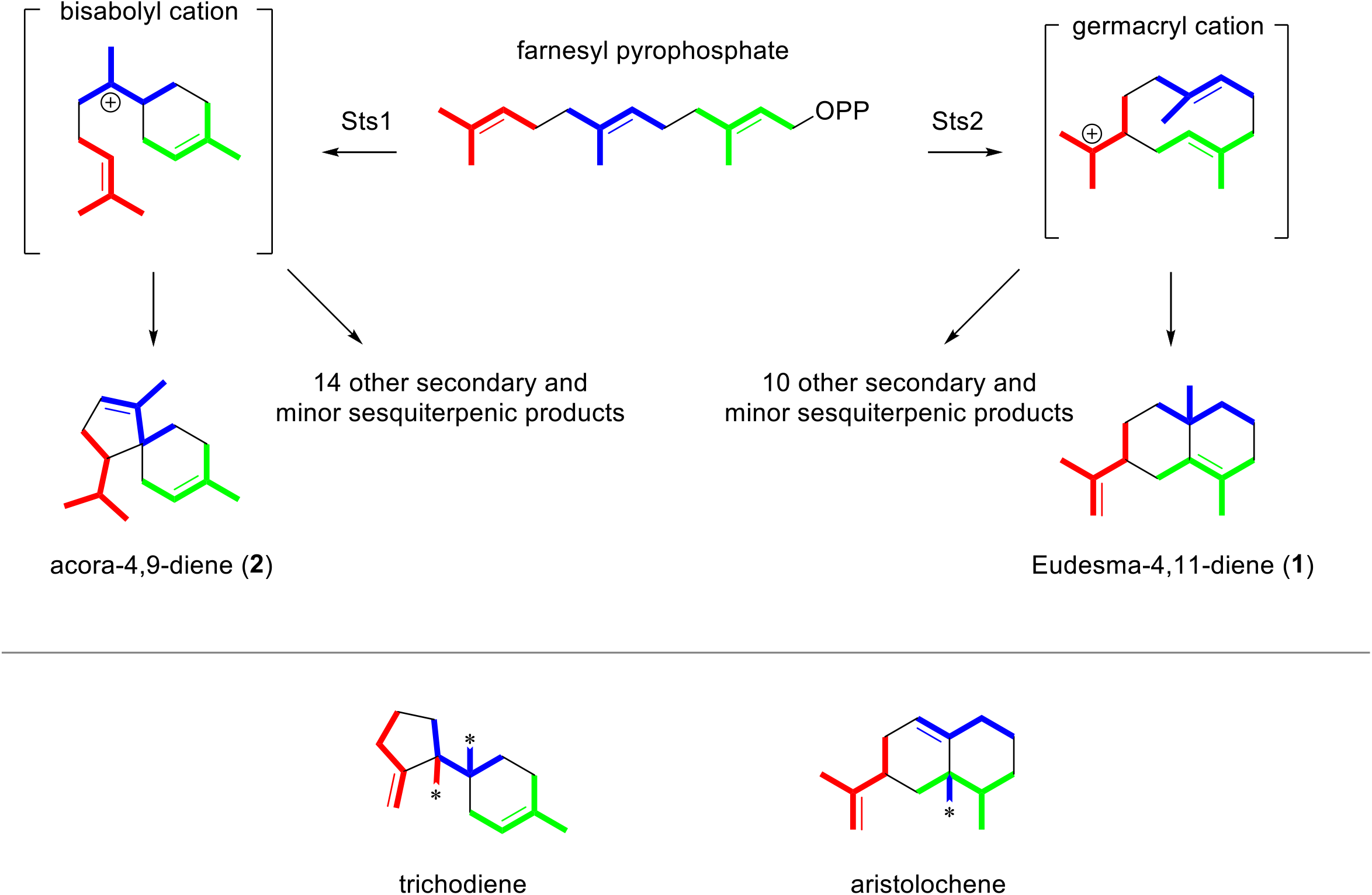
Folding pattern of farnesyl pyrophosphate (FPP) in *P. nodorum* Sts1 and Sts2. Colour indicate the different isoprene units. Asterisks in methyl groups of trichothecene and aristolochene indicate a rearrangement and the colour indicates to which isoprene unit it corresponds.

Even though many biological interactions are established through VOCs, the roles and synthesis of these compounds in fungi are poorly understood. In this study we have found that, in addition to the well-characterised proteinaceous effectors produced by *P. nodorum* as disease determinants, VOCs produced by this pathogen *in vitro* also trigger a response in the host plant as well as having effects on other microorganisms. Additionally, the discovery of *P. nodorum* sesquiterpenes represents the first report of terpenes in this pathogen and complementary techniques were used to link these sesquiterpene structures to their respective biosynthetic genes. Despite these advances, this study exemplifies the many unknowns that remain pertaining to VOCs in fungi and highlights their potential for future research. Unfortunately, the complexity of the volatile bouquets and the chemical signals conveyed by them represent a major challenge when teasing apart biological activities and ecological roles.

## Materials and Methods

### Volatile compound-mediated growth competition assays

To test potential activities of *P. nodorum* VOCs, four fungi were used for growth antagonist assays: *P. nodorum, Fusarium oxysporum* f. sp. *lycopersici, Eutiarosporella tritici-australis* and *Zymoseptoria tritici. Escherichia coli, Pseudomonas syringae, Sinorhizobium meliloti*, and three other bacteria isolated from within surface sterilised wheat seeds, *Bacillus cereus, Sphingobacterium multivorum* and *Flavobacterium* sp. were also used for the growth assays ^73^.

In one side of a segmented Petri dish (9 cm diameter), 25 μl of a *P. nodorum* spore solution (1×10^6^ spores/ml) was inoculated on Fries agar (1.5%) (Table S1) and incubated at 22 °C in 12-12 h dark and light. After two weeks, the other compartment of the segmented Petri dish was inoculated with the test organisms. The fungi were inoculated onto potato dextrose agar (PDA), the bacteria on Lysogeny broth (LB) agar. Water agar (1%) was used for the germination of wheat (Triticum aestivum cv. Grandin) and *Medicago truncatula* (Table S1). The effect of the VOCs was then visually monitored daily.

To assess the phytotoxic and fungistatic effect of 2-methyl-1-propanol, 2-methyl-1-butanol, 3-methyl-1-butanol and 2-phenylethanol, each of the compounds was placed on a 1 cm ^2^ filter paper on a section of a segmented Petri dish ^73^. On the other half of the dish was placed either wheat seeds or *P. nodorum* was inoculated, on the appropriate medium as described above. An additional treatment was also performed containing a mix of these compounds in the proportions found in the chromatographic analysis of *P. nodorum* VOCs. 1 mM of each of the pure compounds (74 to 122 ppm) and 100 ppm for the mix were used in these assays, considering a free internal volume of the petri dish of 48.9 cm^3^ (the total volume minus 15 ml of test medium) ^73^.

### Volatile molecule analysis

To identify the individual components of the VOCs, slanted Fries agar head-space (HS) vials (20 ml) were inoculated with *P. nodorum*. Vials with cotton stoppers were incubated for one week at 22 °C in 12-12 h dark and light cycles. Vials were sealed with silicon/Teflon septa crimp caps 24 hr prior to the analysis. Three mock-inoculated vials were used as controls. To calculate the retention indices, 5 μl of an alkane mix at 20 ppm in CH_2_Cl_2_ was added to a HS vial ^73,74^. To confirm the identity of ethyl acetate, 2-methyl-1-propanol, 2-methyl-1-butanol, 3-methyl-1-butanol and 2-phenylethanol within the fungal VOCs, a mix following the proportions found in the SPME-GC-MS analysis of the fungal cultures (10:14:26:53:11 respectively) was prepared using pure compounds and 1 μl of the mix was added to a HS vial.

To evaluate the *in planta* production of sesquiterpenes, the distal 5 cm of the second leaf of 2-week old wheat seedlings were excised and sprayed with a 1×10^6^ *P. nodorum* spores/ml solution containing 0.02 % tween 20. The cut end of each leaf was embedded in a HS vial containing 2 ml of water agar (1%). Vials were closed with silicon/Teflon septa crimp caps and incubated for 3 days at 22 °C in 12-12 h dark and light cycles. Three mock-inoculated samples were used as controls.

The solid phase micro-extractions in line with a gas chromatography-mass spectrometry (SPME-GC-MS) analyses were performed in an Agilent 7890A gas chromatograph coupled to a single quadrupole Agilent 5975 mass spectrometer using a Gerstel MPS 2XL autosampler. The column for the analyses was a Varian CP9013-1Factor4 5ms 350 °C: 40 m × 250 μm × 0.25 μm. Elution was performed with He flow at 1.5 ml/min and temperature programmed from 40 °C (hold 3 min) to 180 °C at 4 °C /min and then to 220 °C (hold 5 min) at 10 °C/min. The mass spectrometer was operated in the electron ionisation (EI) mode with ionisation energy of 70 eV and scanning the mass range of *m/z* 40-600. Temperatures were set to: GC inlet, 240 °C; GC transfer line, 240 °C; MS source, 200 °C; and quadrupole 250 °C. Volatiles were adsorbed onto a SPME fibre coated with divinylbenzene/carboxen/polydimethylsiloxane (DVB/CAR/PDMS) (1 cm, 23 Ga, 50/30 μm film thickness, Supelco) for 120 min at 30 °C after a 5 min equilibration. The fibre was desorbed in the injector at 240 °C (splitless mode 2 min). The fibres were conditioned by keeping them in the GC injector at 240 °C for 10 minutes.

Data was acquired using MSD ChemStation E.02.01.1177 (© Agilent Technologies, Inc.). Analysis of the data was performed using ChemStation and MS Search (NIST Mass Spectral Search Program [Version 2.0g] for the NIST/EPA/NIH Mass Spectral Library [NIST Standard Reference Database 1A Version NIST 11] build May 19 2011 (© National Institute of Standards and Technology).

### Disruption of *P. nodorum* sesquiterpene synthases

*P. nodorum* sesquiterpene synthases (Sts), SNOG_03562 (*Sts1*) and SNOG_04807 (*Sts2*), were individually disrupted in *P. nodorum* wild type (SN15) by split marker homologous recombination of a phleomycin resistance cassette. 1.5 Kb 5’ and 3’ flanking regions for each gene were amplified from *P. nodorum* genomic DNA (primers in Table S2). The phleomycin resistance gene was amplified as two overlapping amplicons, *Phl* and *Leo*, from the pAN8-1 plasmid (primers in Table S2) ^75^. 5’ flanks were PCR fused to *Leo* while 3’ flanks were PCR fused to *Phl* (primers in Table S2). *P. nodorum* was then transformed by a PEG-protoplast method as previously reported ^76^.

To assess the copy number of the phleomycin cassette in the transformants, qRT-PCR primers were designed for the phleomycin resistance gene; elongation factor 1α, actin and SnToxA primers were used to normalise the data (primers in Table S2). As a phleomycin single copy reference, a *tox3* knock out strain was used ^9^.

### Characterization *P. nodorum* sesquiterpenes

To isolate and characterise the product of Sts1 and Sts2, the coding sequences were cloned into the linearised plasmid backbone (XW55) from YEplac-ADH2p (primers in Table S2)^11,77^. *In vivo* yeast recombination cloning using each gene and XW55 was performed with the Frozen-EZ Yeast Transformation II Kit™ (Zymo Research, Irvine, CA) and competent *Saccharomyces cerevisiae* BJ5464-NpgA according to manufacturer’s protocol. Positive transformants were selected by PCR from colonies grown on synthetic dropout agar lacking uracil.

Medium scale *P. nodorum* fermentations for the isolation of α-cyperone were performed using 8 l of liquid Fries medium inoculated with 4×10^6^ spores. Cultures were incubated in the dark during 10 days at 22 °C and shaking at 120 rpm.

For the isolation of acora-4,9-diene, and eudesma-4,1-diene and β-elemene, transformed *S. cerevisiae* harbouring *Sts1* or *Sts2* were inoculated into in 6 ml synthetic dropout agar lacking uracil (Table S1) for 72 hours at 28 °C 200 rpm. Each of these seed cultures was used to inoculate 5 l YPD broth (Table S1) and incubated 90 hours at 22 °C at 200 rpm ^73^.

Fungal cultures were lyophilised and low to medium polarity compounds were extracted with dichloromethane. Yeast cultures were centrifuged and the cells subjected to acetone extraction. Both, dichloromethane and acetone were evaporated in a rotary evaporator ^73^. The sesquiterpenes from the extracts were isolated by acetonitrile/hexane partition. Sesquiterpenes were purified by SiO_2_ hexane flash chromatography followed by C18 water/acetonitrile flash chromatography ^73^. Purity of terpenes was assessed by GC-MS. Isopentane was used to recover the terpenes from the water-acetonitrile mixture.

GC-MS^2^ was performed to identify β-elemene and α-cyperone by comparison with standards in a Finnigan TraceGC ultra (Thermo Scientific) coupled to an iontrap Finnigan Polaris Q (Thermo Scientific) mass spectrometer. β-elemene was injected onto a BPX70 30m × 0.25mm id (SGE Analytical Science) while α-cyperone in a Varian CP9013-1Factor4 5ms column, which were eluted with He (inlet pressure 15 psi; injection port 200 °C; interface 250 °C; source 200 °C). For β-elemene the column was temperature programmed from 60 °C (hold 1 min) to 100 °C at 25 °C/min, then to 150 at 10 °C/min, and finally to 240 °C at 10 °C/min (hold 3.5 min) ^73^; for α-cyperone the program started at 60 °C (hold 1 min) to 200 °C at 30 °C/min, then to 220 at 3 °C/min, and finally to 325 °C at 30 °C/min (hold 1 min) ^73^. The mass spectrometer was operated in the electronic ionisation (EI) mode with ionisation energy of 70 eV, scanning the mass range of *m/z* 50-450. For MS/MS experiments, the precursor ions were selected with a peak width of 1.0 amu over 12ms. The ions were excited at 1V for 15ms with q = 0.3 and the products scanned over a mass range of *m/z* 100-250. Data analysis was performed employing Xcalibur™ 1.4 (©Thermo Scientific) software.

The eudesma-4,11-diene (**1**) and the acora-4,9-diene (**4**) were dissolved in CDCl_3_ and analysed by NMR. ^1^H NMR, ^13^C NMR, HSQC and HMBC were performed in an Avance™III HD 300 MHz NanoBay NMR device (Bruker) ^73^.

### *P. nodorum* infection assays

To test the requirement of *Sts* genes for infection, the mutants of *P. nodorum* lacking the *Sts1* and *Sts2* genes were inoculated onto the second leaf of two-week-old wheat seedlings (cv. Axe) which was attached to a styrofoam platform using double sided sticky tape and spayed with a spore solution (1×10^6^ spores/ml containing 0.02% tween 20). 0.02% tween 20 was used as control. Inoculated seedlings were incubated for 48h at 22°C in a dark moisture chamber. After the initial 48h incubation, the inoculated plants were grown at 85% humidity, 20 °C during the day and 12 °C at night with 16-8 hr light/dark cycles. The leaves were collected at five days post inoculation leaves to evaluate the disease.

## Acknowledgements

MJMG is a recipient of an Australian Government Endeavour Award and a Mexican CONACYT scholarship. YHC is Australian Research Council Future Fellow (FT160100233).

## Author Contributions

MJMG, YHC and PSS contributed to the design and concept of the study. MJMG, SB, OM, CW and HYYY contributed to the experimentation and collection of data. YHC and RAB contributed to data analysis. MJMG, YHC and PSS wrote the manuscript. All authors intellectually contributed to the study and revision of the manuscript.

## Additional Information

The authors declare no competing interest

## References

1 Oliver, R. P., Friesen, T. L., Faris, J. D. & Solomon, P. S. *Stagonospora nodorum*: From pathology to genomics and host resistance. Annu. Rev. Phytopathol. 50, 23–43 (2012).

2 Friesen, T. L. et al. Emergence of a new disease as a result of interspecific virulence gene transfer. Nat. Genet. 38, 953–956 (2006).

3 Liu, Z. et al. SnTox3 acts in effector triggered susceptibility to induce disease on wheat carrying the Snn3 gene. PLoS Pathog 5, e1000581 (2009).

4 Liu, Z. et al. The cysteine rich necrotrophic effector SnTox1 produced by *Stagonospora nodorum* triggers susceptibility of wheat lines harboring Snn1. PLoS Pathog 8, e1002467 (2012).

5 Breen, S., Williams, S. J., Winterberg, B., Kobe, B. & Solomon, P. S. Wheat PR-1 proteins are targeted by necrotrophic pathogen effector proteins. The Plant Journal 88, 13–25 (2016).

6 Liu, Z. et al. SnTox1, a *Parastagonospora nodorum* necrotrophic effector, is a dual-function protein that facilitates infection while protecting from wheat-produced chitinases. New Phytol. 211, 1052–1064 (2016).

7 McDonald, M. C. & Solomon, P. S. Just the surface: advances in the discovery and characterization of necrotrophic wheat effectors. Curr. Opin. Microbiol. 46, 14–18 (2018).

8 Tan, K.-C. et al. Functional redundancy of necrotrophic effectors – consequences for exploitation for breeding. Frontiers in Plant Science 6 (2015).

9 Tan, K.-C. et al. Sensitivity to three *Parastagonospora nodorum* necrotrophic effectors in current Australian wheat cultivars and the presence of further fungal effectors. Crop and Pasture Science 65, 150–158 (2014).

10 Chooi, Y.-H. et al. An In planta-expressed -polyketide synthase produces (R)-mellein in the wheat pathogen *Parastagonospora nodorum*. Appl. Environ. Microbiol. 81, 177–186 (2015).

11 Chooi, Y.-H., Muria-Gonzalez, M. J., Mead, O. L. & Solomon, P. S. *SnPKS19* encodes the polyketide synthase for alternariol mycotoxin biosynthesis in the wheat pathogen *Parastagonospora nodorum*. Appl. Environ. Microbiol. 81, 5309–5317, doi:10.1128/aem.00278-15 (2015).

12 Chooi, Y.-H. et al. Functional genomics-guided discovery of a light-activated phytotoxin in the wheat pathogen *Parastagonospora nodorum* via pathway activation. Environ. Microbiol. 19, 1975–1986 (2017).

13 Li, H. et al. Chemical Ecogenomics-Guided Discovery of Phytotoxic α-Pyrones from the Fungal Wheat Pathogen *Parastagonospora nodorum*. Organic Letters 20, 6148–6152 (2018).

14 Chooi, Y.-H., Muria-Gonzalez, M. J. & Solomon, P. S. A genome-wide survey of the secondary metabolite biosynthesis genes in the wheat pathogen *Parastagonospora nodorum*. Mycology 5, 192–206 (2014).

15 Muria-Gonzalez, M. J., Chooi, Y.-H., Breen, S. & Solomon, P. S. The past, present and future of secondary metabolite research in the Dothideomycetes. Mol. Plant Pathol. 16, 92–107 (2015).

16 Kesselmeier, J. & Staudt, M. Biogenic volatile organic compounds (VOC): an overview on emission, physiology and ecology. J. Atmos. Chem. 33, 23–88 (1999).

17 Kanchiswamy, C. N., Malnoy, M. & Maffei, M. E. Chemical diversity of microbial volatiles and their potential for plant growth and productivity. Frontiers in Plant Science 6 (2015).

18 Ipcho, S. V. S. et al. Transcriptome analysis of *Stagonospora nodorum*: gene models, effectors, metabolism and pantothenate dispensability. Mol. Plant Pathol. 13, 531–545 (2012).

19 Chen, Y.-J. & Lin, W.-Y. Total synthesis of (±)-acoradiene via radical cyclization. Tetrahedron 49, 10263–10270 (1993).

20 Iwabuchi, H., Yoshikura, M. & Kamisako, W. Studies on the Sequiterpenoids of Panax ginseng C. A. MEYER. III. Chem. Pharm. Bull. 37, 509–510 (1989).

21 Chitarra, G. S., Abee, T., Rombouts, F. M., Posthumus, M. A. & Dijksterhuis, J. Germination of *Penicillium paneum* Conidia Is Regulated by 1-Octen-3-ol, a Volatile Self-Inhibitor. Appl. Environ. Microbiol. 70, 2823–2829 (2004).

22 Polizzi, V. et al. Autoregulatory properties of (+)-thujopsene and influence of environmental conditions on its production by *Penicillium decumbens*. Microb. Ecol. 62, 838–852 (2011).

23 Papaleo, M. C. et al. Sponge-associated microbial Antarctic communities exhibiting antimicrobial activity against *Burkholderia cepacia* complex bacteria. Biotechnol. Adv. 30, 272–293 (2012).

24 Romoli, R. et al. GC–MS volatolomic approach to study the antimicrobial activity of the antarctic bacterium *Pseudoalteromonas* sp. TB41. Metabolomics 10, 42–51 (2013).

25 Tirranen, L. S. & Gitelson, I. I. The role of volatile metabolites in microbial communities of the LSS higher plant link. Adv. Space Res. 38, 1227–1232 (2006).

26 Fialho, M. B., Moraes, M. H. D. d., Tremocoldi, A. R. & Pascholati, S. F. Potential of antimicrobial volatile organic compounds to control *Sclerotinia sclerotiorum* in bean seeds. Pesqu. Agropecu. Bras. 46, 137–142 (2011).

27 Hua, S. S., Beck, J. J., Sarreal, S. B. & Gee, W. The major volatile compound 2-phenylethanol from the biocontrol yeast, *Pichia anomala*, inhibits growth and expression of aflatoxin biosynthetic genes of *Aspergillus flavus*. Mycotoxin Res. 30, 71–78 (2014).

28 Lester, G. Inhibition of growth, synthesis, and permeability in *Neurospora crassa* by phenethyl alcohol. J. Bacteriol. 90, 29–37 (1965).

29 Liu, P. et al. Mechanisms of action for 2-phenylethanol isolated from *Kloeckera apiculata* in control of *Penicillium molds* of citrus fruits. BMC Microbiol. 14, 1–14 (2014).

30 Chang, P. K., Hua, S. S. T., Sarreal, S. B. L. & Li, R. W. Suppression of aflatoxin biosynthesis in *Aspergillus flavus* by 2-phenylethanol is associated with stimulated growth and decreased degradation of branched-chain amino acids. Toxins 7, 3887–3902 (2015).

31 Splivallo, R., Novero, M., Bertea, C. M., Bossi, S. & Bonfante, P. Truffle volatiles inhibit growth and induce an oxidative burst in *Arabidopsis thaliana*. New Phytol. 175, 417–424, doi:10.1111/j.1469-8137.2007.02141.x (2007).

32 Farag, M. A., Ryu, C.-M., Sumner, L. W. & Paré, P. W. GC–MS SPME profiling of rhizobacterial volatiles reveals prospective inducers of growth promotion and induced systemic resistance in plants. Phytochemistry 67, 2262–2268 (2006).

33 Ryu, C.-M. et al. Bacterial volatiles promote growth in *Arabidopsis*. Proc. Natl. Acad. Sci. U. S. A. 100, 4927–4932 (2003).

34 McNeal, K. S. & Herbert, B. E. Volatile organic metabolites as indicators of soil microbial activity and community composition shifts. Soil Sci. Soc. Am. J. 73 (2009).

35 Pritchard, L. & Birch, P. A systems biology perspective on plant–microbe interactions: Biochemical and structural targets of pathogen effectors. Plant Sci. (Amsterdam, Neth.) 180, 584–603 (2011).

36 Baldwin, I. T., Halitschke, R., Paschold, A., von Dahl, C. C. & Preston, C. A. Volatile Signaling in Plant-Plant Interactions: “Talking Trees” in the Genomics Era. Science 311, 812–815 (2006).

37 Caruso, C. M. & Parachnowitsch, A. L. Do Plants Eavesdrop on Floral Scent Signals? Trends Plant Sci. 21, 9–15 (2016).

38 Cha, D. H. et al. Eavesdropping on Plant Volatiles by a Specialist Moth: Significance of Ratio and Concentration. PLoS One 6, e17033 (2011).

39 Chu, S. S., Jiang, G. H. & Liu, Z. L. Insecticidal compounds from the essential oil of Chinese medicinal herb *Atractylodes chinensis*. Pest Manag. Sci. 67, 1253–1257 (2011).

40 Lee, T. K. & Vairappan, C. S. Antioxidant, antibacterial and cytotoxic activities of essential oils and ethanol extracts of selected South East Asian herbs. J. Med. Plants Res. 5, 5284–5290 (2011).

41 Takahashi, M., Inouye, S. & Abe, S. *Anti-Candida* and radical scavenging activities of essential oils and oleoresins of *Zingiber officinale* Roscoe and essential oils of other plants belonging to the family Zingiberaceae. Drug Discoveries Ther. 5, 238–245 (2011).

42 Wang, J., Li, J., Li, S., Freitag, C. & Morrell, J. J. Antifungal activities of *Cunninghamia lanceolata* heartwood extractives. BioResources 6, 606–614 (2011).

43 Yoshida, T., Endo, K., Ito, S. & Nozoe, T. Chemical constituents in the essential oil of the leaves of *Chamaecyparis taiwanensis*. Yakugaku Zasshi 87, 434–439 (1967).

44 Ayoub, N., Schultze, W. & Lass, D. Volatile constituents of the medicinal fungus chaga Inonotus obliquus (Pers.: Fr.) Pilát (Aphyllophoromycetideae). 11, 55–60 (2009).

45 Brock, N. L. & Dickschat, J. S. PR toxin biosynthesis in *Penicillium roqueforti*. ChemBioChem 14, 1189–1193 (2013).

46 Rabe, P., Citron, C. A. & Dickschat, J. S. Volatile terpenes from actinomycetes: a biosynthetic study correlating chemical analyses to genome data. ChemBioChem 14, 2345–2354 (2013).

47 Rösecke, J., Pietsch, M. & König, W. A. Volatile constituents of wood-rotting basidiomycetes. Phytochemistry 54, 747–750 (2000).

48 Yuan, Z., Chen, Y., Mao, L., Zhang, C. & Chen, L. Activity of three volatile-producing endophytic *Muscodor* strains in controlling fruit postharvest disease and analysis of the volatile components. Linye Kexue 49, 83–89 (2013).

49 Braekman, J. C., Remacle, A. & Roisin, Y. Soldier defensive secretion of three Amitermes species. Biochem. Syst. Ecol. 21, 661–666 (1993).

50 Krasulova, J. et al. Chemistry and anatomy of the frontal gland in soldiers of the sand termite *Psammotermes hybostoma*. J. Chem. Ecol. 38, 557–565 (2012).

51 Naya, Y., Miyamoto, F. & Takemoto, T. Formation of enantiomeric sesquiterpenes in the secretions of scale insects. Experientia 34, 984–986 (1978).

52 Adio, A. M. (−)-trans-β-elemene and related compounds: occurrence, synthesis, and anticancer activity. Tetrahedron 65, 5145–5159 (2009).

53 Baker, R. et al. Chemical composition of the frontal gland secretion of *Syntermes* soldiers (Isoptera, Termitidae). J. Chem. Ecol. 7, 135–145 (1981).

54 Bowers, W. S., Nishino, C., Montgomery, M. E., Nault, L. R. & Nielson, M. W. Sesquiterpene progenitor, germacrene A: an alarm pheromone in aphids. Science 196, 680–681 (1977).

55 Everaerts, C., Roisin, Y., Le Quere, J. L., Bonnard, O. & Pasteels, J. M. Sesquiterpenes in the frontal gland secretions of nasute soldier termites from New Guinea. J. Chem. Ecol. 19, 2865–2879 (1993).

56 Fassotte, B. et al. First evidence of a volatile sex pheromone in lady beetles. PLoS One 9, e115011 (2014).

57 Omura, H., Honda, K. & Feeny, P. From terpenoids to aliphatic acids: further evidence for late-instar switch in osmeterial defense as a characteristic trait of swallowtail butterflies in the tribe papilionini. J. Chem. Ecol. 32, 1999–2012 (2006).

58 Quintana, A. et al. Interspecific variation in terpenoid composition of defensive secretions of European *Reticulitermes* termites. J. Chem. Ecol. 29, 639–652 (2003).

59 Fischer, G., Schwalbe, R., Möller, M., Ostrowski, R. & Dott, W. Species-specific production of microbial volatile organic compounds (MVOC) by airborne fungi from a compost facility. Chemosphere 39, 795–810, doi:http://dx.doi.org/10.1016/S0045-6535(99)00015-6 (1999).

60 Jeleń, H. H. Volatile sesquiterpene hydrocarbons characteristic for *Penicillium roqueforti* strains producing PR toxin. J. Agric. Food Chem. 50, 6569–6574 (2002).

61 Sanchez-Fernandez, R. E. et al. Antifungal volatile organic compounds from the endophyte *Nodulisporium* sp. strain GS4d2II1a: a qualitative change in the intraspecific and interspecific interactions with *Pythium aphanidermatum*. Microb. Ecol. 71, 347–364 (2016).

62 Bradfield, A. E., Hegde, B. H., Rao, B. S., Simonsen, J. L. & Gillam, A. E. 152. α-Cyperone, a sesquiterpene ketone from the oil of *Cyperus rotundus*. Journal of the Chemical Society (Resumed), 667–677 (1936).

63 Dadang Ohsawa, K., Kato, S. & Yamamoto, I. Insecticidal compound in tuber of *Cyperus rotundus* L. against the diamondback moth larvae. J. Pestic. Sci. 21, 444–446 (1996).

64 Komai, K., Iwamura, J. & Ueki, K. Isolation, identification and physiological activities of sesquiterpenes in purple nutsedge tubers. J. Weed Sci. Technol. 22, 14–18 (1977).

65 Mojab, F., Vahidi, H., Nickavar, B. & Kamali-nejad, M. Chemical components of essential oil and antimicrobial effects of rhizomes from *Cyperus rotundus* L. J. Med. Plants 8, 91–97+186 (2009).

66 Pirzada, A. M. et al. *Cyperus rotundus* L.: traditional uses, phytochemistry, and pharmacological activities. J. Ethnopharmacol. 174, 540–560 (2015).

67 Krohn, K. et al. Bioactive natural products from the endophytic fungus *Ascochyta* sp. from *Meliotus dentatus* – Configurational assignment by solid-state CD and TDDFT calculations. Eur. J. Org. Chem. 2007, 1123–1129 (2007).

68 Hashidoko, Y., Tahara, S. & Mizutani, J. in Z.Naturforsch.(C) Vol. 47 353 (1992).

69 Kaiser, R. & Naegeli, P. Biogenetically significant components in vetiver oil. Tetrahedron Lett. 13, 2009–2012 (1972).

70 Mazzoni, V., Tomi, F. & Casanova, J. A daucane-type sesquiterpene from *Daucus carota* seed oil. Flavour Fragr. J. 14, 268–272 (1999).

71 Cane, D. E. Enzymic formation of sesquiterpenes. Chem. Rev. 90, 1089–1103 (1990).

72 Christianson, D. W. Unearthing the roots of the terpenome. Curr. Opin. Chem. Biol. 12, 141–150 (2008).

73 Muria-Gonzalez, M. J. Secondary metabolites in the Parastagonospora nodorum-wheat pathosystem. Doctor of Philosophy (PhD) thesis, The Australian National University.

74 Kováts, E. Gas-chromatographische charakterisierung organischer verbindungen. Teil 1: retentionsindices aliphatischer halogenide, alkohole, aldehyde und ketone. Helv. Chim. Acta 41, 1915–1932 (1958).

75 van Den Dool, H. & Dec. Kratz, P. A generalization of the retention index system including linear temperature programmed gas—liquid partition chromatography. J. Chromatogr. A 11, 463–471 (1963).

76 Mattern, I. E., Punt, P. J. & Van den Hondel, C. A. M. J. J. A vector for *Aspergillus* transformation conferring phleomycin resistance. Fungal Genetics Reports 35, 1 (1988).

77 Solomon, P. S., Tan, K.-C., Sanchez, P., Cooper, R. M. & Oliver, R. P. The Disruption of a Gα Subunit Sheds New Light on the Pathogenicity of *Stagonospora nodorum* on Wheat. Mol. Plant-Microbe Interact. 17, 456–466 (2004).

78 Lee, K. M. & DaSilva, N. A. Evaluation of the *Saccharomyces cerevisiae* ADH2 promoter for protein synthesis. Yeast 22, 431–440 (2005).

