## Supplementary Data for "Volatile molecules secreted by the wheat pathogen *Parastagonospora nodorum* are involved in development and phytotoxicity"

### Contents

**Table S1**. Media recipes ... 2

**Table S2**. List of primers ... 3

**Figure S1.** Organisms that showed low or no sensitivity to *P. nodorum* VOCs. ... 4

**Figure S2.** GC-MS from the isolated sesquiterpenes produced by *STS01* expressed in yeast. . 5

**Figure S3.** GC-MS from the isolated sesquiterpenes produced by *STS02* expressed in yeast. . 6

**Figure S4.** MSMS Identification of β-elemene. ... 7

**Figure S5.** MSMS Identification of α-cyperone. ... 8

**Figure S6.** Eudesma-4,11-diene (1) H NMR (300MHz, CDCl3) ... 9

**Figure S7**. Eudesma-4,11-diene (**1**) C NMR (300MHz, $CDCl_3$) ... 10

**Figure S8**. Eudesma-4,11-diene (**1**) HSQC (300MHz, $CDCl_3$) ... 11

**Figure S9**. Eudesma-4,11-diene (**1**) HMBC (300MHz, $CDCl_3$) ... 12

**Figure S10**. Acora-4,9-diene (**4**) H NMR (300MHz, $CDCl_3$) ... 13

**Figure S11**. Acora-4,9-diene (**4**) C NMR (300MHz, $CDCl_3$) ... 14

**Figure S12**. Acora-4,9-diene (**4**) HSQC (300MHz, $CDCl_3$) ... 15

**Figure S13**. Acora-4,9-diene (**4**) HMBC (300MHz, $CDCl_3$) ... 16

**Table S3.** NMR data for eudesma-4,11-diene (**1**) (300MHz, $CDCl_3$). ... 17

**Table S4.** NMR data for acora-4,9-diene (**4**) (300MHz, $CDCl_3$) ... 18

**Figure S14.** Pathogenicity test of sts1 and sts2 on wheat 5 days post inoculation. ... 19

**Figure S15**. Phylogenetic tree of fungal terpene synthases. ... 20

**Table S1**. Media recipes

| **Media** | **Composition per litre of media** |
|---|---|
| Water agar (1%) | 10 g Agar |
| Potato Dextrose Agar (PDA) | PDA: 39 g Difco$^{TM}$ dehydrated PDA |
| Fries (pH 6) | 30.0 g sucrose<br>5.0 g yeast extract<br>5.0 g ammonium tartrate<br>1.0 g $NH_4NO_3$<br>1.0 g $KH_2PO_4$<br>0.5 g $MgSO_4{\cdot}7H_2O$<br>0.13 g CaCl<br>0.1 g NaCl<br>If solid media add 15.0 g Agar |
| Lysogeny Broth (LB) | 10.0 g NaCl<br>10.0 g triptone<br>5.0 g yeast extract<br>If solid media add 15.0 g Agar |
| YPD | 10 g yeast extract<br>20.0 g triptone<br>After autoclaving add:<br>20.0 g glucose |
| Synthetic dropout media lacking uracil (pH 5.8) | 6.7 g Yeast base media (no amino acids)<br>1.6 g amino acid mix lacking uracil<br>After autoclaving add 30 g glucose<br>If solid media add 15.0 g Bacto Agar |

**Table S2**. List of primers

| Name | forward | reverse |
|---|---|---|
| Phl | ccctcgttgaccaagaatc | aagttcgtggacacgacct |
| Leo | gacgtgaccctgttcatca | gagcattcactaggcaacca |
| Sts01_5flank | ttctggttcggaataagctctt | gagcattcactaggcaa-atttattgtgcgcgttcg |
| Sts01_3flank | ctaagaaccagttgctccc-tctttgcacctacgatttttcc | cattcctgattcctcttgtttg |
| Sts02_5flank | gcgcggaatggtgtaaagt | gagcattcactaggcaa-aaagatgatctcgcctgcc |
| Sts02_3flank | ctaagaaccagttgctcccc-tgaaccggcgtaccagttta | accaacacgtcagtgacgac |
| XW_STS01 | atcaactatcaactattaactatatcgtaatacca-atgtcccattcccgccacga | tgtcatttaaattagtgatggtgatggtgatgcac-cacctccacggcagtgacct |
| XW_STS02_exon1 | atcaactatcaactattaactatatcgtaatacca-atggctccgatagatgcatca | gacccatggagaagcacatgactccggacatga agcctttaccaagatcggcatgtc |
| XW_STS02_exon2 | tactttgagtaccgacatgccgatcttggtaaa ggcttcatgtccggagtcatgtg | tgtcatttaaattagtgatggtgatggtgatgcac-accccaaatccttccttcattg |
| Phleo_qPCR | ggaagttcgtggacacga | gacgtgaccctgttcatca |
| Elng.f_qPCR | atgatcgacgtttccacc | tcaatagcctcgaggagg |
| Actin_qPCR | agtcgaagcgtggtatcct | acttggggttgatgggag |
| SnToxA_qPCR | gtccgtctgtcaacaacatcg | tcagttcccacgagcctatagc |

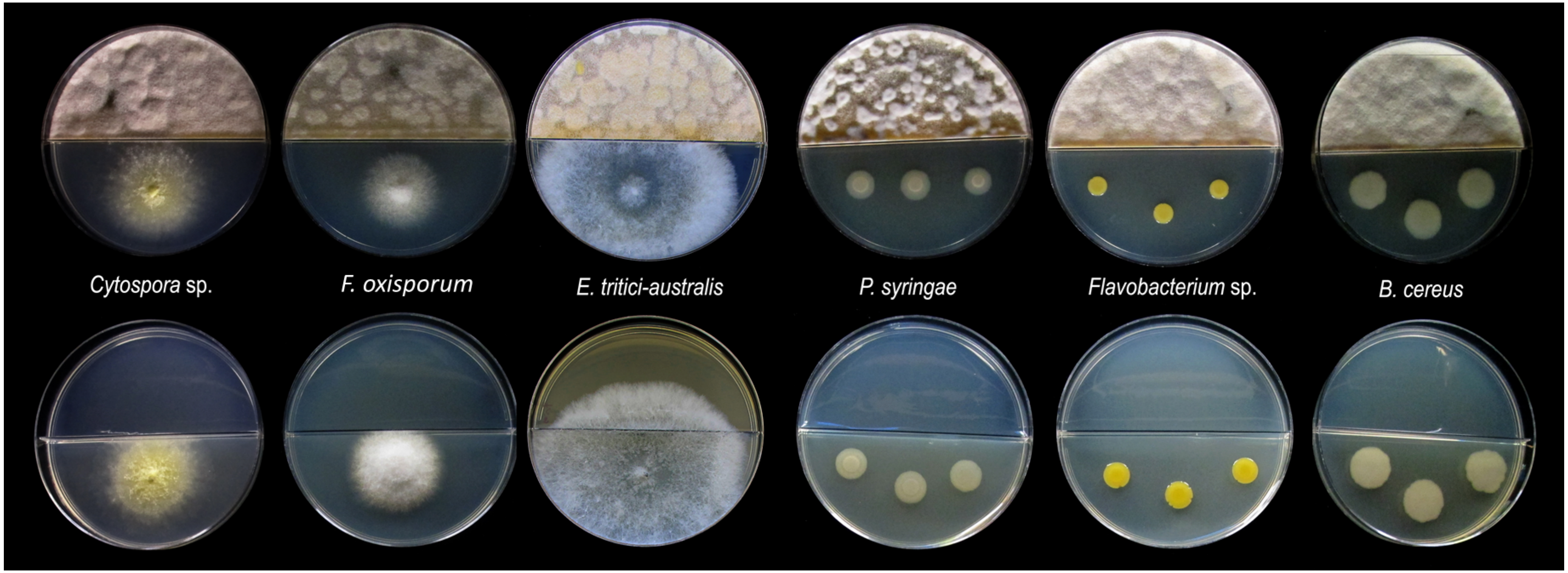


**Figure S1.** Organisms that showed low or no sensitivity to *P. nodorum* VOCs emitted *in vitro*.
Split plate assay of the effect of VOCs produced by wild type P. nodorum (SN15). The top row of plates show effect (or lack of effect) caused by SN15 VOCs over *Cytospora sp., Fusarium oxisporum, Eutiarosporella tritici-australis, Pseudomonads syringae, flavobacterium sp.* and *Bacillus cereus*. The bottom row of plates presents each test organism next to the control, media only: non-inoculated Fries agar.

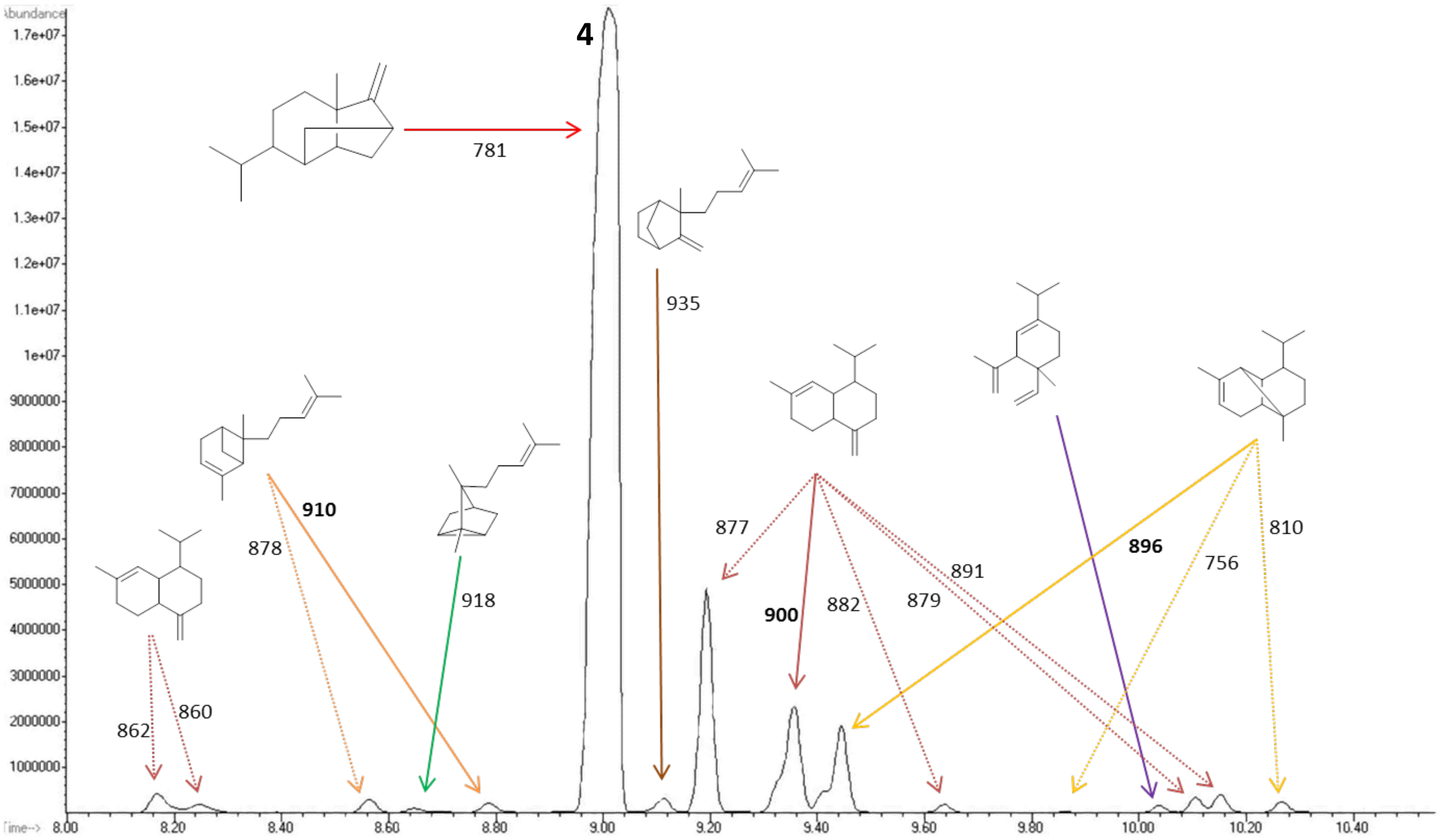


**Figure S2.** GC-MS from the isolated sesquiterpenes produced by *STS01* expressed in yeast**.**

The peak corresponding to **4** is marked with its number. The presented structures correspond to the best match within the NIST library. The numbers are the score given by the NIST library (the score could go up to 1000). The proposed structure for **4** is isosativene but did not show a good spectral match to the NIST library as the lower score indicates.

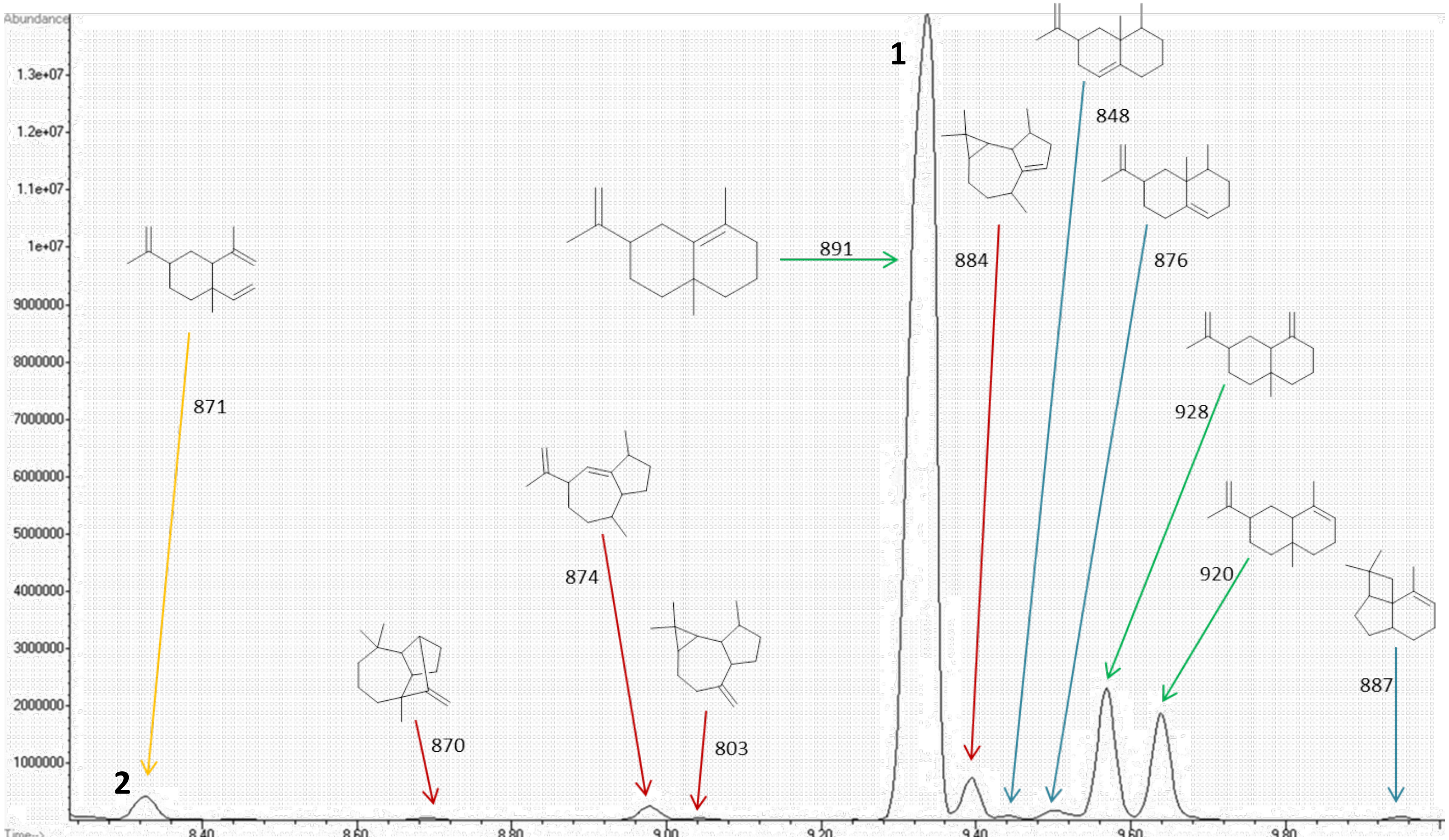


**Figure S3.** GC-MS from the isolated sesquiterpenes produced by *STS02* expressed in yeast**.**
The peaks corresponding to **1** and **2** are marked with their number. The presented structures correspond to the best match within the NIST library. The numbers are the score given by the NIST library (the score could go up to 1000).

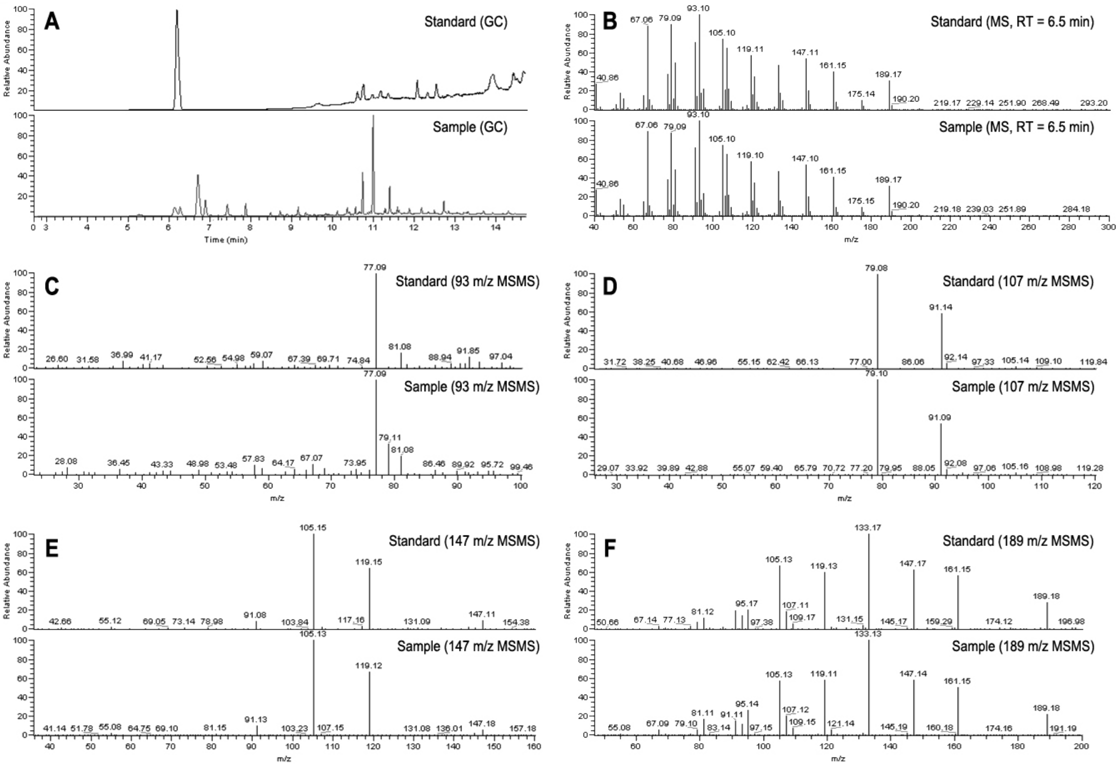


**Figure S4.** MSMS Identification of β-elemene.
Figure shows the MS chromatogram (A), MS spectrum (B) and $MS^2$ spectra of ion $93^+$ (C), $107^+$ (D), $147^+$ (E) and $189^+$ (F). All panels compare acquired data from the spectra (above), and the *P. nodorum* sample (below).

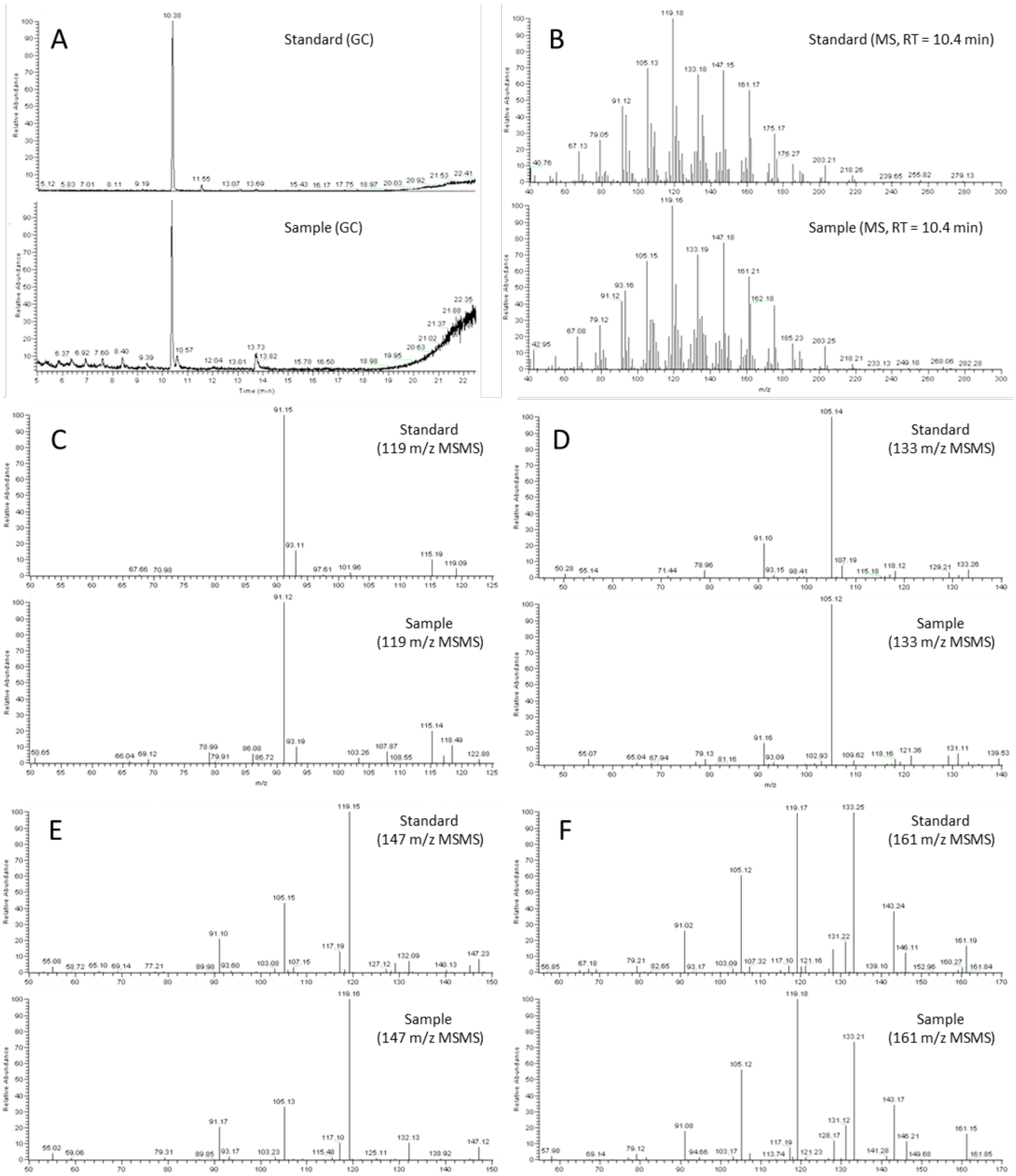


**Figure S5.** MSMS Identification of α-cyperone.
Figure shows the MS chromatogram (A), MS spectrum (B) and $MS^2$ spectra of ion $119^+$ (C), $133^+$ (D), $147^+$ (E) and $161^+$ (F). All panels compare acquired data from the spectra (above), and the purified sample from *P. nodorum* (below).

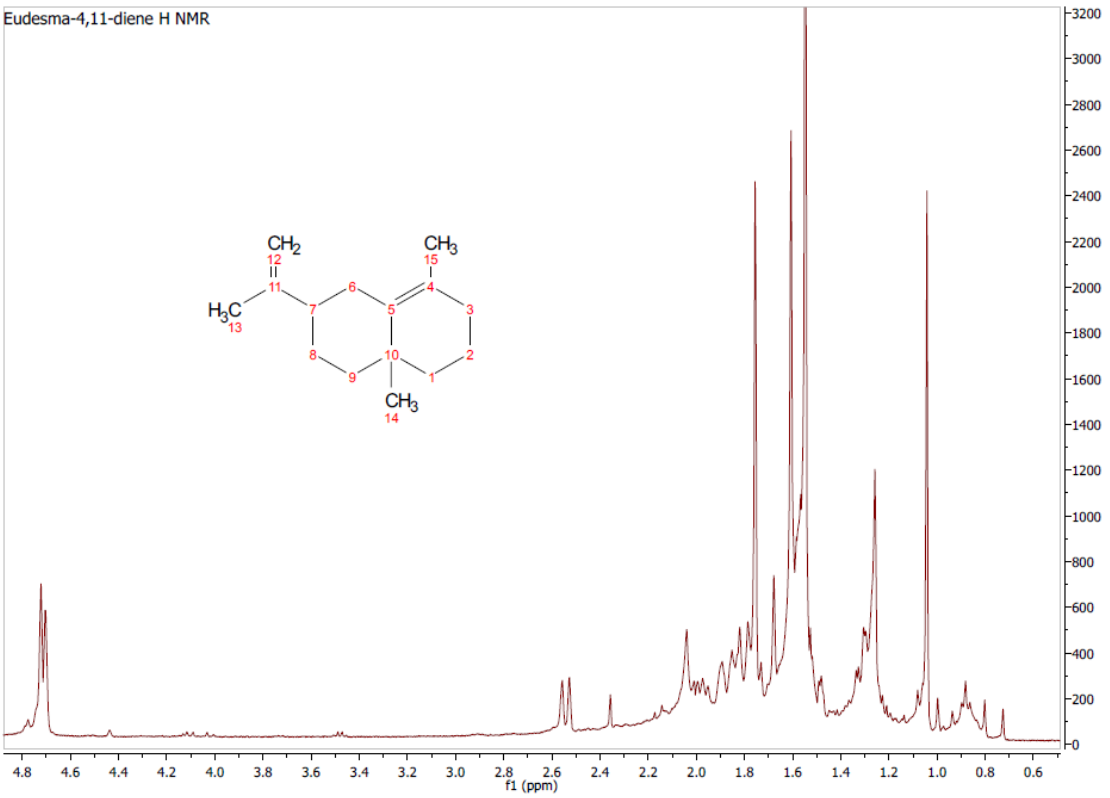


**Figure S6.** Eudesma-4,11-diene (1) H NMR (300MHz, CDCl3)

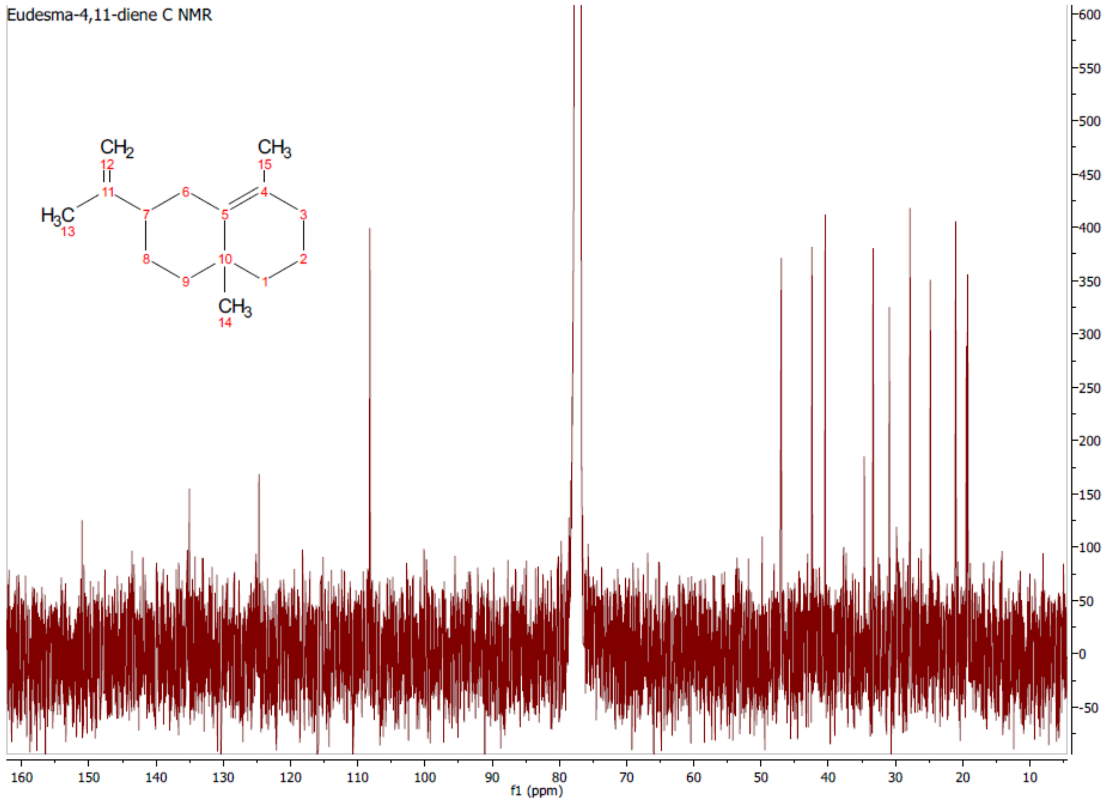


**Figure S7**. Eudesma-4,11-diene (**1**) C NMR (300MHz, $CDCl_3$)

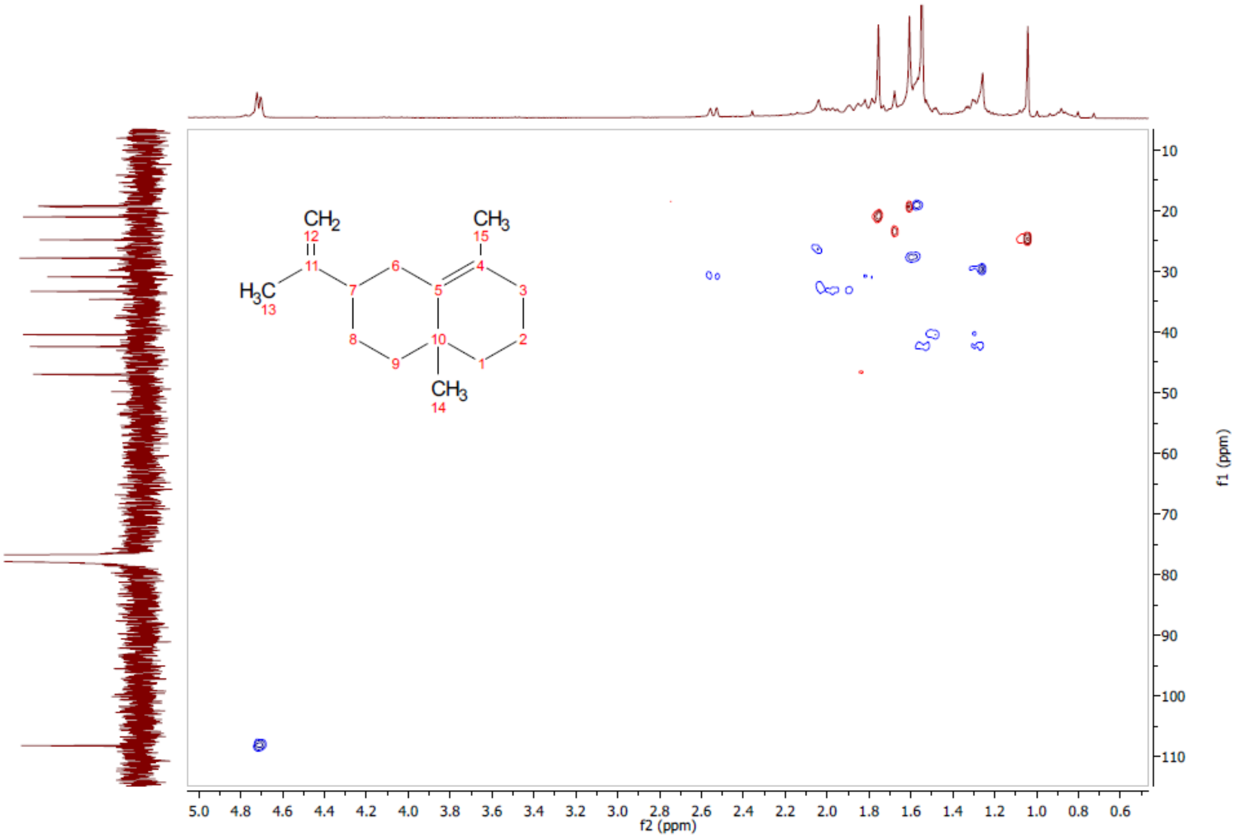


**Figure S8**. Eudesma-4,11-diene (**1**) HSQC (300MHz, $CDCl_3$)

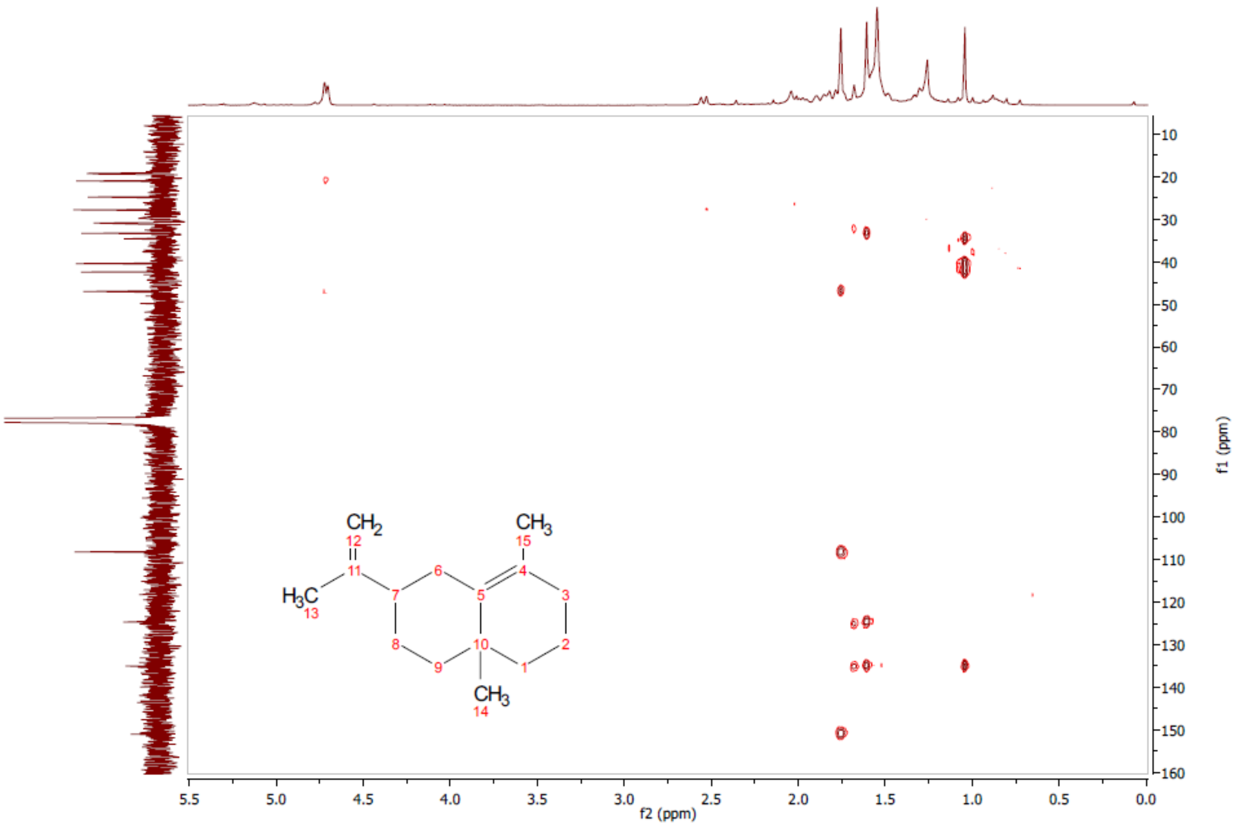


**Figure S9**. Eudesma-4,11-diene (**1**) HMBC (300MHz, $CDCl_3$)

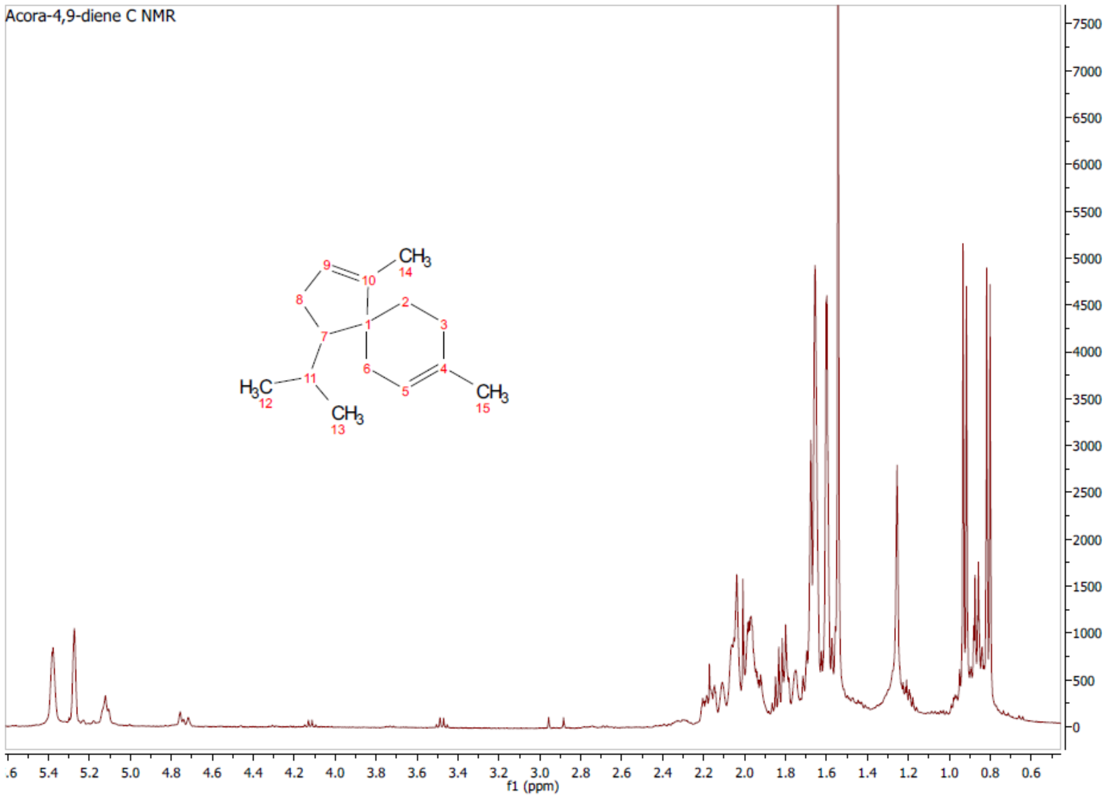


**Figure S10**. Acora-4,9-diene (**4**) H NMR (300MHz, $CDCl_3$)

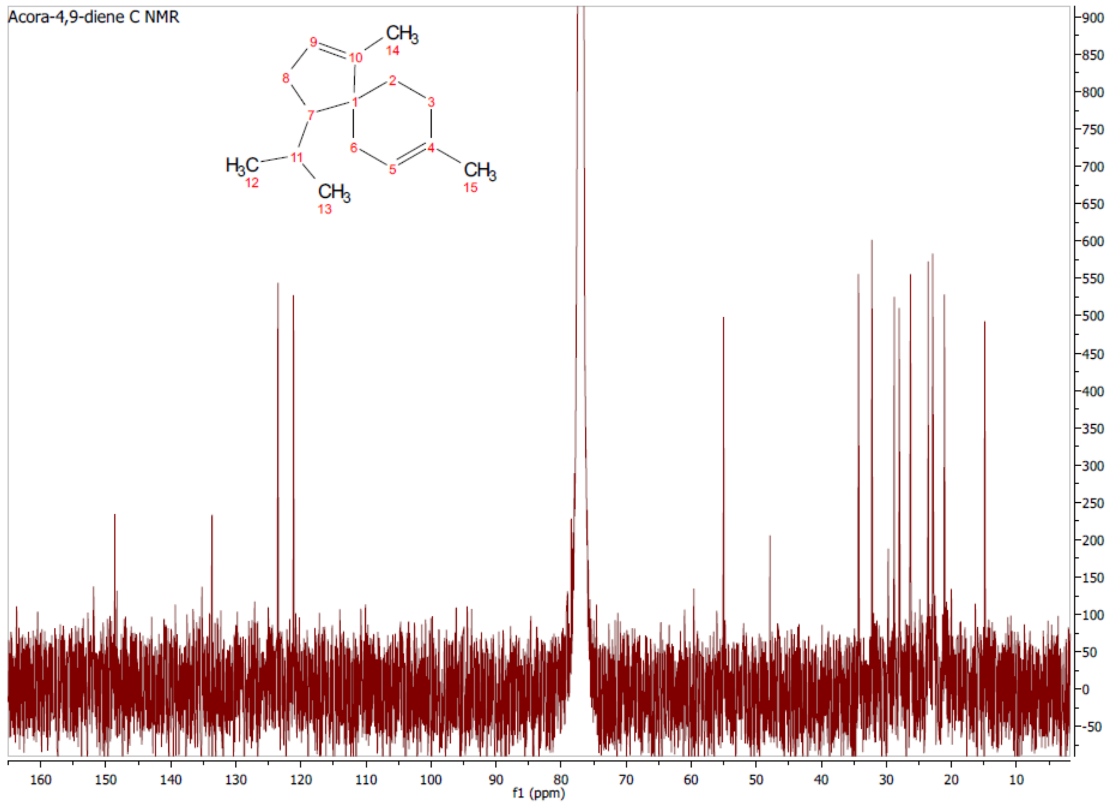


**Figure S11**. Acora-4,9-diene (**4**) C NMR (300MHz, $CDCl_3$)

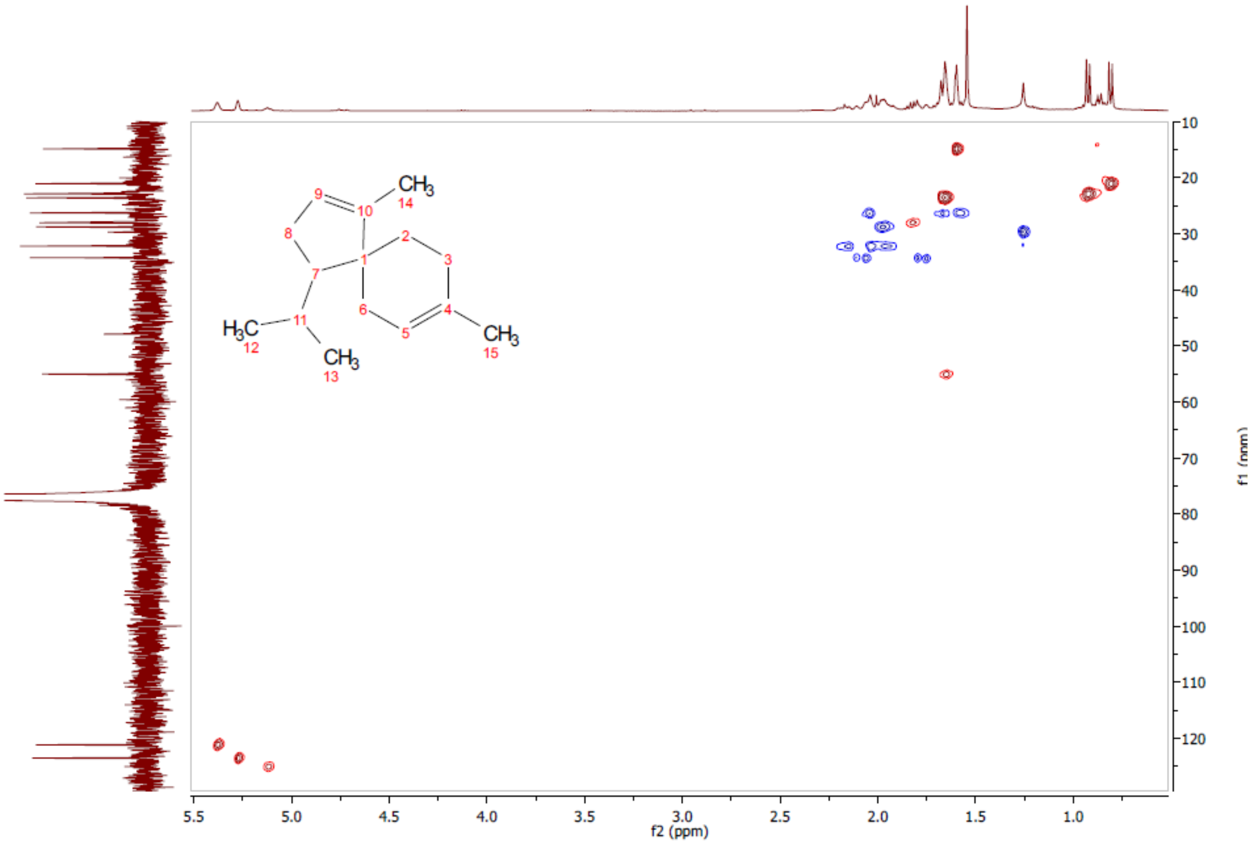


**Figure S12**. Acora-4,9-diene (**4**) HSQC (300MHz, $CDCl_3$)

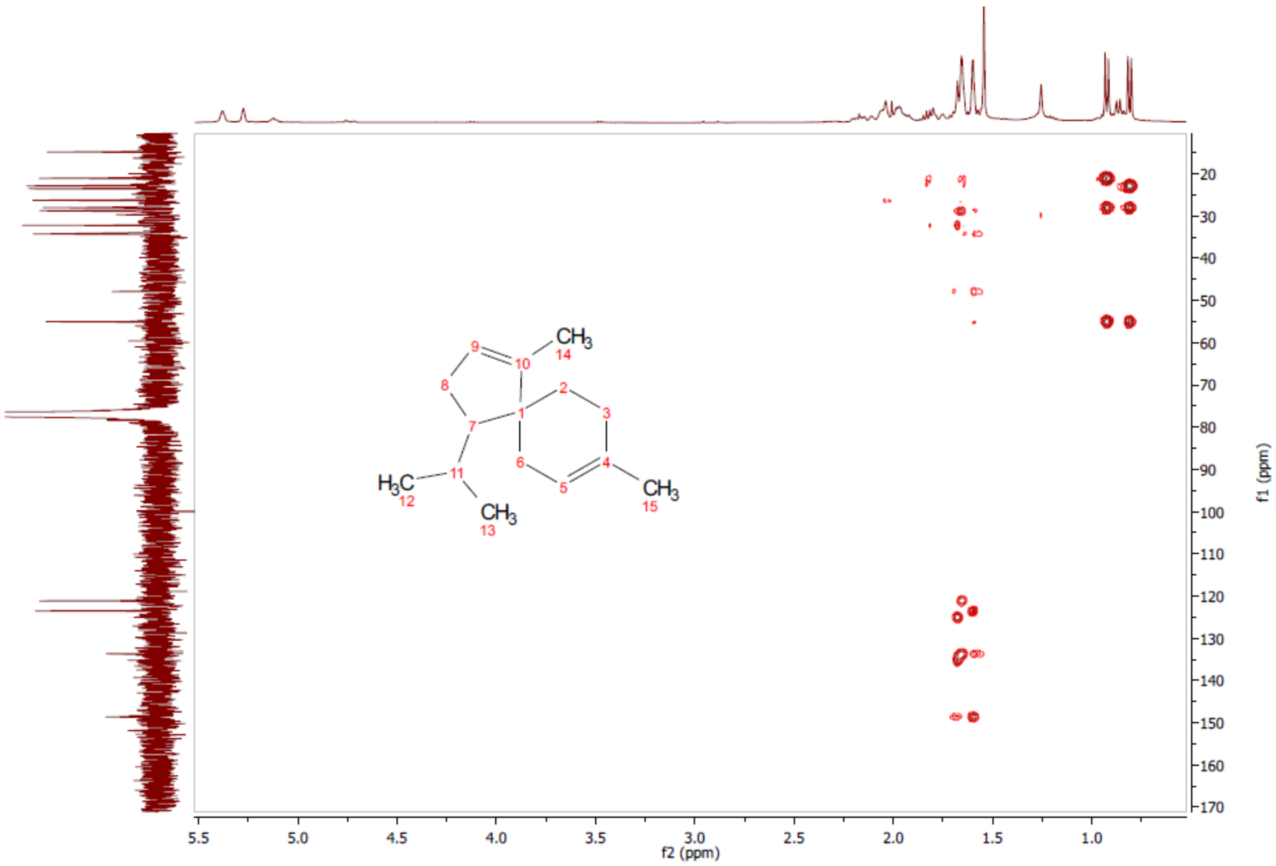


**Figure S13**. Acora-4,9-diene (**4**) HMBC (300MHz, $CDCl_3$)

**Table S3.** NMR data for eudesma-4,11-diene (**1**) (300MHz, $CDCl_3$).
Chemical shifts (δ) for C ($C^{13}$ NMR) and H ($H^1$ NMR) and C→H correlation at 2-4 bonds distance (HMBC) are presented.

| Carbon number | C δ (ppm) | Hδ$_a$ (ppm) | Hδ$_b$ (ppm) | HMBC C →H |
|---|---|---|---|---|
| 1[#] | 40.45 | 1.32 | 1.5 | 14 |
| 2 | 19.26 | 1.57 | | |
| 3 | 33.32 | 1.89 | 1.99 | 15 |
| 4 | 124.69 | - | - | 15 |
| 5 | 135.04 | - | - | 14, 15 |
| 6 | 30.91 | 2.54 | | |
| 7 | 46.98 | 1.87 | - | 13 |
| 8 | 27.83 | 1.59 | | |
| 9[#] | 42.44 | 1.29 | 1.51 | 14 |
| 10 | 34.64 | - | - | 14 |
| 11 | 151.01 | - | - | 13 |
| 12 | 108.19 | 4.70 | 4.72 | 13 |
| 13 | 21.03 | 1.75 | - | 12 |
| 14 | 24.8 | 1.04 | - | |
| 15 | 19.44 | 1.61 | - | |

[#] Data no enough to distinguish between C1 and C9

**Table S4.** NMR data for acora-4,9-diene (**4**) (300MHz, $CDCl_3$).
Chemical shifts (δ) for C ($C^{13}$ NMR) and H ($H^1$ NMR) and C→H correlation at 2-4 bonds distance (HMBC) are presented.

| Carbon number | C δ (ppm) | H$\delta_a$ (ppm) | H$\delta_b$ (ppm) | HMBC C →H |
|---|---|---|---|---|
| 1 | 47.94 | - | - | 2a, 2b$^+$ |
| 2 | 26.78 | 1.58* | 1.67 | 3b |
| 3 | 34.77 | 1.78 | 2.09 | 2a |
| 4 | 149.09 | - | - | 2b$^+$, 15 |
| 5 | 123.97 | 5.27 | - | 15 |
| 6 | 29.22 | 1.98* | | 2a, 2b$^+$, 14$^+$ |
| 7 | 28.4 | 1.83 | - | 12, 13 |
| 8 | 32.63 | 1.97 | 2.18 | 7, 14$^+$ |
| 9 | 121.6 | 5.38 | - | 14$^+$ |
| 10 | 134.4 | - | - | 2a, 2b$^+$, 14$^+$ |
| 11 | 55.49 | 1.65* | - | 12, 13 |
| 12 | 21.48 | 0.81 | - | 7, 11$^+$, 13 |
| 13 | 23.29 | 0.93 | - | 12 |
| 14 | 23.99 | 1.66* | - | |
| 15 | 15.29 | 1.6* | - | |

*Signals not clear on H NMR, values obtained from HSQC
$^+$HMBC correlations with H 2b, H11 and H14 are temptative because singnals are too close

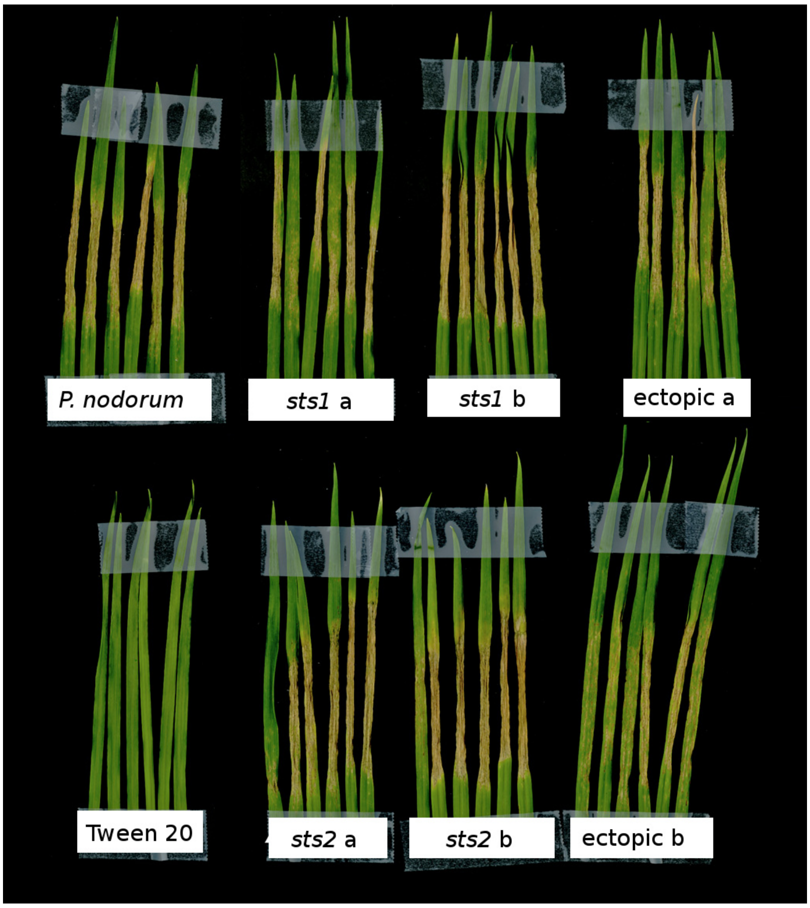


**Figure S14.** Pathogenicity test of sts1 and sts2 on wheat 5 days post inoculation. Two sts1, two sts2 strains and two ectopic mutants were inoculated on wheat cv. Axe. SN15 was used as a positive control and 0.02% tween 20 solution as negative control.

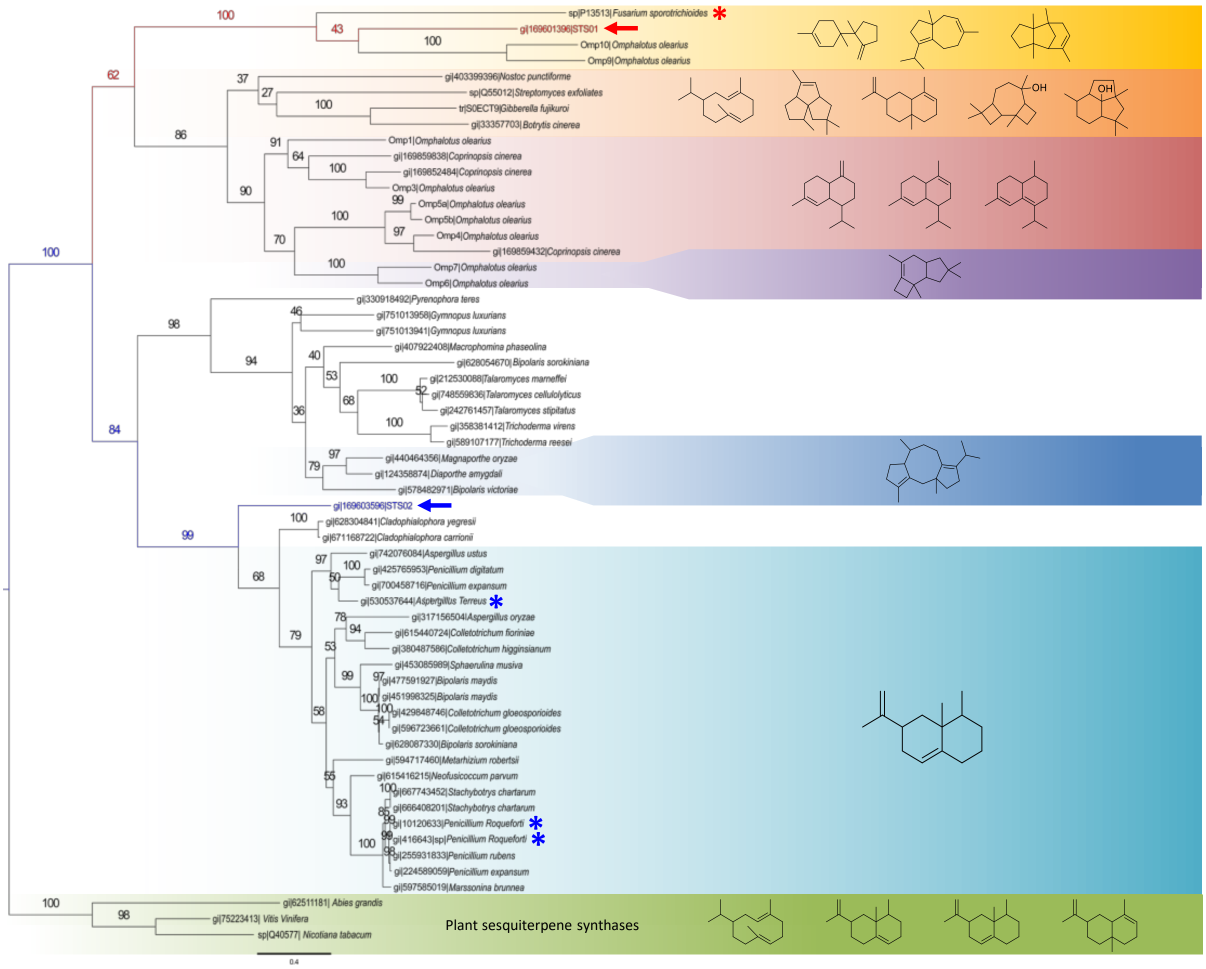


**Figure S15**. Neighbour joining phylogenetic tree of fungal terpene synthases (plant sesquiterpene synthases as outliers). Red arrow indicates *P. nodorum* Sts1, blue arrow indicates *P. nodorum* Sts2, red asterisk indicates a characterised trichodiene synthase, and blue asterisks indicate characterised aristolochene synthases. The presented chemical structures correspond to some of the reported products of the terpene synthases on each group.
